# Regenerative Grazing as a Climate Change Mitigation Strategy: A Systematic Review

**DOI:** 10.1101/2025.10.11.677897

**Authors:** Docker B. Clark, Shelby C. McClelland, Jasmine A. Dillon, Matthew N. Hayek

**Affiliations:** New York University; New York University, Stony Brook University; Cornell University

**Author notes:** Corresponding Author - Matthew N. Hayek.

**Keywords:** regenerative agriculture, livestock, climate mitigation, SOC, grazing, carbon sequestration

## Abstract

Ruminant livestock production is a meaningful contributor to global greenhouse gas (GHG) emissions. Regenerative grazing has been proposed as a climate change mitigation strategy for offsetting livestock GHG emissions by potentially causing additional soil organic carbon (SOC) sequestration. We review the current state of the literature and collect reported and inferred SOC sequestration from regenerative grazing experiments, ranking their study designs by ascending evidentiary strength: observational, cross-sectional and longitudinal. Across 28 studies that compare regenerative to conventional grazing, the reported and inferred SOC sequestration rates in regenerative grazing treatments varied widely, with the highest values reported in lower-strength observational studies. Among higher-strength cross-sectional studies and longitudinal studies, median additional SOC sequestration rates in regenerative grazing treatments were not significantly different than zero, indicating no significant enhancement of SOC across all regenerative grazing interventions. Currently, higher-strength evidence does not support widespread claims of SOC sequestration from regenerative grazing. To clarify its potential for climate mitigation, a greater number of longitudinal studies are needed.

## 1. Introduction

Ruminant meat and dairy production account for 9% to 13% of global anthropogenic greenhouse gas (GHG) emissions (FAO, 2022; Halpern et al., 2022). One notable GHG mitigation strategy has sparked popular imagination: “regenerative grazing”, which may result in soil organic carbon (SOC) sequestration. Regenerative grazing lacks an agreed-upon definition, but encompasses livestock management practices aimed at improving soil and ecosystem health, farm profit, and resiliency (Dillon and Machmuller, 2021). These practices often involve rotating herds across smaller subdivided pastures at high stocking densities to mimic natural grazing.

Highly-visible public presentations have asserted that regenerative grazing meaningfully offsets cattle emissions, or even offset emissions from other economic sectors (Leu 2024).

Recent scientific assessments reach diverging conclusions about SOC sequestration from regenerative grazing (Eshel et al., 2025; Pelton et al., 2024). However, these studies derived estimates from only a few studies. Additionally, these reviews have not analyzed experimental designs, i.e. whether they were potentially prone to confounding variables.

Our systematic review assembles and evaluates evidence addressing the question: does regenerative grazing induce SOC sequestration? Specifically, what is the quantity and quality of evidence that SOC sequestration occurred and was additional to counterfactual conditions, e.g. under conventional grazing? We assemble SOC sequestration outcomes from all regeneratively-managed ruminant livestock grazing experiments to date, and evaluate SOC results based on a hierarchy of evidence based on their experimental designs.

## 2. Methods

### 2.1. Grazing management definitions

To facilitate comparison between regenerative and conventional grazing, we define “conventional grazing” as methods that allow animals to access pastures for the entirety of the grazing season without providing rest. This is also commonly referred to as “continuous grazing,” “traditional grazing,” and “set-stocking”. We define “regenerative grazing” as a suite of methods that subdivide larger pastures into “paddocks” that are alternatively intensively grazed by the entire herd or subdivisions of it, then rested intermittently with periods of no grazing. This rotational approach is a frequent but not universal tenet of regenerative agriculture, a term that lacks a consensus definition (Giller et al., 2021). Many terms in scientific and agricultural parlance meet our functional definition, including: adaptive multi-paddock grazing, mob grazing, management intensive grazing, and holistic planned management.

### 2.2. Literature review

We conducted a systematic review of the literature following PRISMA guidelines (Fig. S1), for results quantifying SOC outcomes of regenerative grazing. To compile these articles, we conducted series of database searches from 2002 extending to the present (accessed: Feb 5, 2025) using Web of Science and Google Scholar. We searched Web Of Science for all studies containing a wide array of search terms related to regenerative grazing and its synonyms, plus soil carbon outcomes (Box S1). This query returned 86 studies. We included studies that A) compare regenerative grazing and/or conventional grazing management; B) report soil organic matter (SOM%, SOC%, SOC stocks, or SOC flux; C) present original in-situ soil carbon measurements; and D) report the duration of time pastures were under either grazing treatment. A total of 23 of the 86 Web Of Science retrieved studies met our initial criteria.

Additional Google Scholar searches returned 1 unique study. Searching references from literature reviews appearing in the initial search resulted in 4 more studies that met our criteria. Lastly, references cited by the regenerative grazing SOC analyses that met our search criteria were carefully examined for inclusion, but none met all of our criteria. In total, we found 28 unique studies meeting our inclusion criteria. Several studies (N = 11) reported data using a management type falling outside of our definition of conventional grazing as a counterfactual for regenerative grazing management. These data are accounted for in our auxiliary analysis but were not suitable for our main analysis. In addition to SOC, we catalogued the studies’ location, species, animal stocking densities and rates, soil sampling depth(s), and land use history.

### 2.3. Categorization of study designs

We categorized the 28 studies that met our criteria into four experimental design categories according to a hierarchy of evidentiary strength: (1) Time-Series Observational studies, which reported SOC from regeneratively grazed sites, using either true time series or chronosequence measurements, but without an observed analogous conventionally grazed site; (2) Comparative Observational studies, that compared adjacent sites that differed in one or more biotic or abiotic confounding variables (e.g., soil texture, climate, land use history) and reported only a single SOC measurement; (3) Matched Quasi-Experimental (MQE) Cross-Sectional studies, which directly compare adjacent regenerative and conventional sites that are otherwise similar with respect to all reported biotic and abiotic properties the authors deemed likely to affect SOC, but only report SOC at a single time point, and; (4) MQE Longitudinal studies reporting at least two SOC measurements over a period of ≥1 year. MQE designs were ranked higher than comparative observational studies because they attempt to control for confounding variables including land use history and different starting baseline SOC quantities. No true experimental studies, which would randomize grazing treatments among sites, were found in our literature search.

### 2.4. SOC stocks

We converted all studies’ SOC stock data into common units of metric tons of carbon per hectare (MgC ha^-1^) which some studies report directly (N = 16). In cases, soil carbon concentration units were reported (%SOC or SOC g/kg) and we used the following equation to convert to C stock:

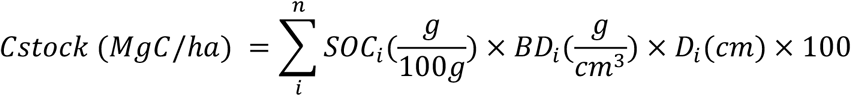

where BD = bulk density of a soil sample at depth i, D = the depth in centimeters of each stratum of soil, SOC = soil organic carbon concentration at depth i, and 100 is the conversion factor of gC cm^-2^ to MgC ha^-1^. When soil C concentration was reported in %SOM, we first converted SOM to SOC using a 0.58 conversion factor (McClelland et al., 2021; Lunt, 1931; Broadbent, 1953; Howard, 1965). In cases where we were limited by a lack of BD measurements by depth, we applied a single value to all depths: (1) Where only one BD was reported (N = 1), we applied this to all depths; (2) Where none were reported (N = 3), an estimated value from literature (USDA, 2019) was applied to all depth strata based on soil textures reported within their respective studies.

### 2.5. SOC fluxes

Annual fluxes from these studies (MgC ha^-1^ y^-1^) were either reported directly, or we calculated them according to the common method within other studies in their evidentiary category. All Time-Series Observational studies reported their own SOC flux rates which we incorporated directly. However, for the comparative observational studies that did not quantify flux, we took the difference in total reported SOC stock between treatment and control sites divided by the duration of the treatment.

For cross-sectional studies, differences in C stock between treatment and control sites were divided by the number of years since the site was given contrasting grazing treatments. If treatment durations differed, we calculated the mean of the durations. Some studies did not explicitly organize adjacent regenerative and conventional pasture sites into pairs. We paired these sites according to similar animal stocking densities/rates. If no stocking information was available, SOC was averaged across treatment sites and compared to the average across control sites.

For MQE Longitudinal studies, fluxes were calculated by taking a difference of differences: the difference in SOC stock from the first (baseline) and last (most recent) sampling year, then the difference between paired regenerative and conventional sites, dividing the result by number of years since first and last measurements.

When fluxes were reported directly we collected them from their parent study regardless of how the authors calculation methods differed from those we outline above.

### 2.6. Statistical analysis

All statistical analysis was conducted using R version 4.3.2 (R Core Team, 2023). Data manipulation and visualizations were facilitated by use of the “tidyverse”, “_matrixStats”, and “_ggrepel” packages (Wickham et al. 2019; Bengtsson, 2025; Slowikowski 2024). Means and medians for each category were calculated with 95% confidence intervals using bootstrapping by randomly resampling SOC fluxes over N = 1001 synthetic vectors with replacement (Efron, 1979).

## 3. Results

We compiled 28 regenerative grazing studies in which SOC fluxes were either reported or could be inferred using methods common in the literature. Combined, these studies represent 49 pasture sites (one pair of co-located continuous vs. regenerative case-control pastures = one pasture site). In most studies, pastures were grazed by cattle (N = 16), while others were grazed by sheep (N = 5), or had mixed cattle with sheep and/or goats (N = 5). The Americas and Australia were disproportionately represented: 93% of studies, i.e. all but two. In descending order: 46% of studies or 57% of sites were in North America; 32% of studies or 27% of sites were in Australia; 14% of studies and 10% of sites were in South America. Two remaining studies, with a combined three sites, concerned sheep grazing in China and Spain.

Time-series observational studies show the greatest maximum values for annual SOC sequestration. Median and mean SOC sequestration rates were 3.59 (2.29 - 7.10 95% CI)

MgC ha^-1^ y^-1^ and 4.33 (2.29 – 7.10) MgC ha^-1^ y^-1^, respectively, both of which were significantly greater than zero (Table 1). Comparative observational studies’ median SOC sequestration was 0.33 MgC ha^-1^ y^-1^ and was not significantly greater than zero; the mean was significantly greater than zero at 2.42 (0.02 – 4.93) MgC ha^-1^ y^-1^, but overlaps zero when studies with insufficient soil depth data are excluded (−0.43 – 6.61) MgC ha^-1^ y^-1^.

**Table 1:** Summary statistics within each study design category of SOC sequestration in MgC ha^-1^ y^-1^ with bootstrapped 95% confidence intervals (positive numbers = sequestration). Columns on the right were calculated from data excluding studies with depth measurements that were deemed insufficient.

|  | All studies |  |  |  | Studies with sparse depth data excluded |  |  |  |
| --- | --- | --- | --- | --- | --- | --- | --- | --- |
| Study design | Studies | SOC sequestration |  |  | Studies | SOC sequestration |  |  |
|  |  | Range | Median (95% CI) | Mean (95% CI) |  | Range | Median (95% CI) | Mean (95% CI) |
| Time-Series Observational | 3 | 2.29 – 7.1 | 3.59 (2.29 – 7.1) | 4.33 (2.29 – 7.1) | 3 | 2.29 – 7.1 | 3.59 (2.29 – 7.1) | 4.33 (2.29 – 7.1) |
| Comparative Observational | 5 | -0.43 – 6.61 | 0.33 (-0.43 – 6.61) | 2.42 (0.02 – 4.93) | 3 | -0.43 – 6.61 | 5.25 (-0.43 – 6.61) | 3.81 (-0.43 – 6.61) |
| MQE Cross-Sectional | 16 | -0.58 – 3.37 | 0.09 (-0.05 – 0.42) | 0.5 (0.06 – 1.05) | 13 | -0.58 – 3.37 | 0.12 (-0.05 – 0.49) | 0.62 (0.06 – 1.24) |
| MQE Longitudinal | 5 | -0.3 – 0.89 | -0.03 (-0.3 – 0.89) | 0.13 (-0.18 – 0.52) | 2 | -0.3 – -0.13 | -0.21 (-0.3 – -0.13) | -0.21 (-0.3 – -0.13) |

The largest category by study count was MQE Cross-Sectional (N = 16). Median SOC sequestration was 0.09 MgC ha^-1^ y^-1^ and was not significantly greater than zero (−0.05 – 0.42). This lack of statistical significance remained when SOC data from studies with sparse depth measurements were excluded (Table 1). In both cases, however, mean SOC sequestration in the MQE cross-sectional study category was significantly greater than zero (Table 1) due to a non-symmetrical distribution of SOC outcomes in this category.

MQE longitudinal studies (N = 5) showed the smallest range of SOC fluxes, and median sequestration of −0.03 MgC ha^-1^ y^-1^, which was not significantly less than zero (−0.03 – 0.89), nor was the mean (Table 1). Three out of the five longitudinal studies measured surface soil only (10 cm).

The same tendencies in medians and means across study categories were observed when we excluded studies with sparse soil depth data (Table 1), when we treated sites as replicates rather than studies (Fig S2), or when we normalized SOC numbers to a common depth (Figs, S3 & S4).

To evaluate farmed livestock species as a potentially confounding variable (Fig S5), we calculated summary statistics for our dataset separating categories by grazed species (Table S1). We evaluated the significance of SOC sequestration on cattle-grazed vs all other grazed ruminant categories (Mixed Large and Small + Small Ruminants) using a Mann-Whitney U, which returned a p-value of 0.0879, i.e. not significant at the 95% confidence level. We performed this test on only studies within the MQE cross-sectional category as this provided the largest and most even sample size of cattle-grazing (N = 9) and non-cattle ruminant grazing (N = 6) studies.

The 2019 case-controlled study conducted by Alemu et al. (2019) collected SOC measurements over time, and provides sufficient data to calculate results in both a cross-sectional and longitudinal manner (Fig 1). We incorporated both, to highlight the difference in apparent SOC sequestration rates from MQE Cross-Sectional versus MQE Longitudinal study designs.

**Fig. 1:**
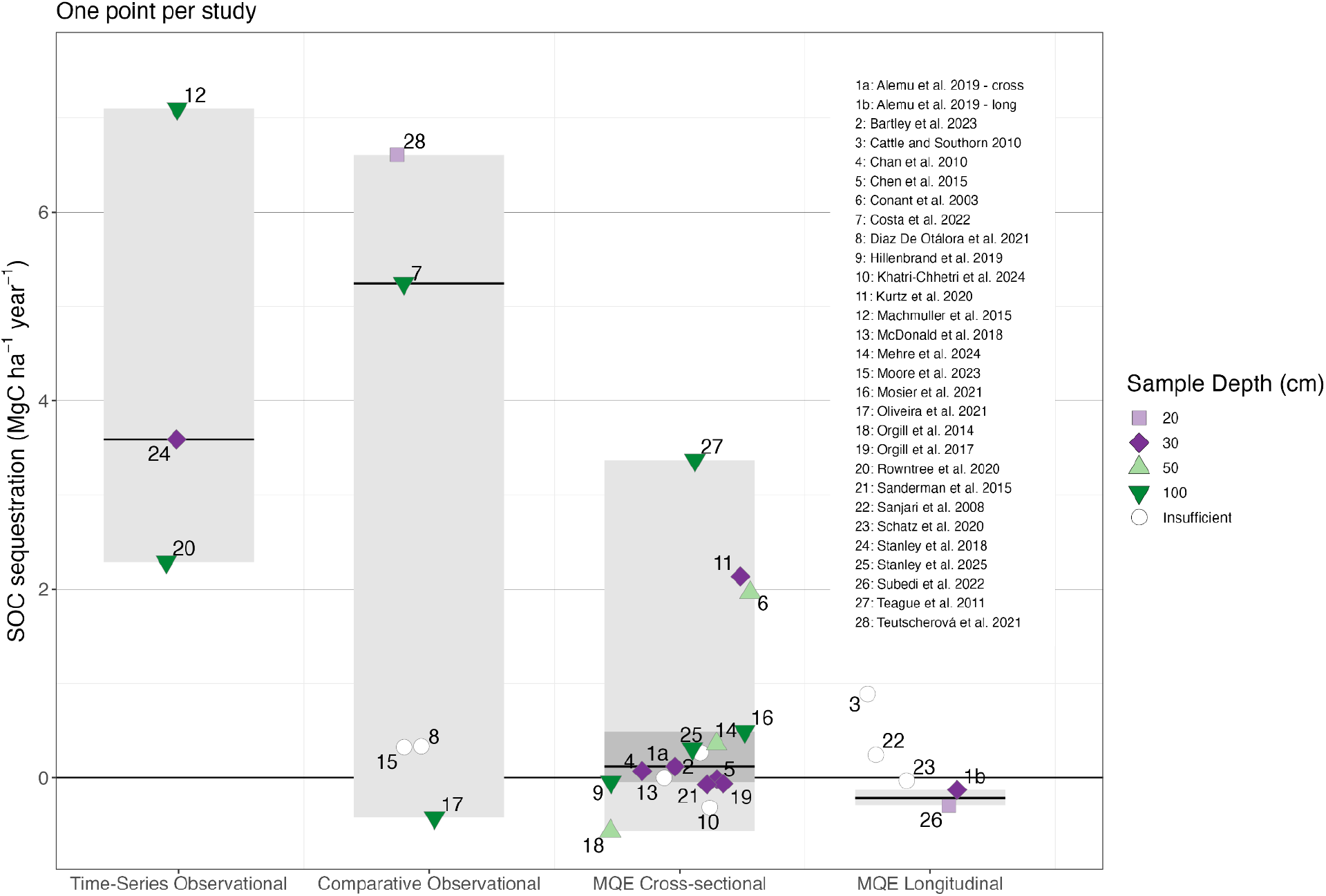
Mean SOC sequestration rates in each regenerative grazing study (points and numbers) and ranges (boxes). Point colors represent the maximum depth at which SOC measurements were conducted. Studies with depth measurements of <20 cm or with only one depth stratum were classified as having insufficient depth data (white points). Light gray boxes represent full ranges of data and black line segments are medians for each category, excluding studies with insufficient depth data. Dark gray box represents interquartile range for MQE Cross sectional category.

## 4. Discussion

### 4.1. Carbon sequestration outcomes from regenerative grazing practices

Experimental designs produced different diverging conclusions about regenerative management’s influence on SOC. The highest-strength studies (MQE longitudinal) did not demonstrate consistent evidence for regenerative grazing enhancing SOC sequestration. This may be due to their small number (N = 5). Only two MQE longitudinal studies had suitable depth measurements, neither of which found an additive effect of regenerative grazing upon SOC over time. For the remaining studies, extrapolating their surface SOC measurements to the entire soil profile would be specious because SOC can move throughout the soil column without resulting in net SOC loss or gain (Hao et al. 2017).

Among MQE Cross-Sectional studies we found a positive *mean* SOC enhancement in regenerative sites, which could suggest that regenerative grazing enhances SOC sequestration. However, there are two important caveats to this finding. First, *median* SOC sequestration was not significantly different from zero, meaning studies were approximately equally likely to find

SOC results in either direction. Second, without repeated measurements (i.e., longitudinal studies) it is not possible to attribute SOC responses to regenerative grazing per se; measured differences in SOC could be an artifact of different baselines from the start of the regenerative grazing experiment.

Despite notable limitations in the evidentiary strength of multiple studies’ design, studies commonly included language that suggested SOC sequestration and/or enhancement was caused by regenerative grazing, e.g. “AMP grazing can contribute to climate change mitigation through SOC sequestration” (Stanley et al. 2018) or “…with significant [SOC] accumulation in the improved grassland area” (Costa et al. 2022), across N = 12 studies. These claims overstate the conclusions warranted by the experimental design, because measured differences may be due to coincidental differences in baseline SOC, land management history, or other confounding environmental variables that the study designs do not control for.

In all studies, differences in SOC could also arise from interventional or observational biases. No studies included in our analysis used a randomized design. This introduces a potential site selection bias if ranchers chose more productive pastures to try novel or risky management methods, or the opposite.

### 4.2. Future recommendations

To substantiate claims about regenerative grazing enhancing or sequestering SOC, more longitudinal studies are needed. Improvements to study design are readily feasible. Among MQE Cross-Sectional study sites, additional work could be done to turn some into Longitudinal studies; if management methods at these sites are presently the same as they were during reported baseline measurements, a single additional set of soil measurements could be sufficient. Ideally, multiple more years of measurement should occur at every site to demonstrate that any net sequestration is not due to an anomalous year of climatic or ecological conditions. Furthermore, additional studies should begin with randomized assignment to control for site selection bias.

Observational studies are especially low-strength evidence that regenerative grazing results in SOC sequestration because land use history is a confounding variable. While all three Time-Series Observational studies reported positive SOC sequestration, their sites all had histories of annual cropping (Machmuller et al. 2015; Stanley et al. 2018; Rowntree et al. 2020). Transitioning to *any* system with perennial roots (e.g. conventional pasture, mowed meadow, orchard, wetland, savanna, or forest) will result in SOC sequestration after tillage and other soil disturbance are removed (Yang et al. 2020). By contrast, MQE Longitudinal designs control for this history. Studies in this category show that additional SOC sequestration from regenerative grazing is closer to and overlaps zero, with the only two studies that took sufficient depth measurements (multiple and greater then 10 cm) showing SOC loss over time (Fig. 1; Table 1). Despite this, observational studies used causal language suggesting that regenerative grazing itself induced SOC sequestration.

We use conventional continuous grazing as a counterfactual for regenerative grazing because it represents a directly comparable land use with the same fundamental goal of livestock production. Other non-grazing land management practices—such as grazing exclusion, hay harvesting, or wild grazing—were present in the literature (N = 11). These counterfactuals were included in our auxiliary analysis (Figs. S6 & S7) because they are qualitatively different from a production standpoint. Relaxed rotational grazing, defined by a lower frequency and/or stocking rate than typical regenerative practices, was also deemed an inappropriate counterfactual for our main analysis and was placed in the auxiliary analysis as it does not represent a distinct alternative land use. When comparing regenerative grazing to these non-continuous grazing counterfactuals, results were mixed. Additional SOC sequestration rates from regenerative grazing still overlapped zero (Figs S6 & S7).

We found no significant difference in soil organic carbon (SOC) between sites grazed by cattle and those grazed by other ruminants (p = 0.09). This indicates that livestock species may not be a major confounding factor in our assessment. Although studies on cattle grazing appeared to show slightly higher average SOC sequestration from regenerative sites, the limited sample size for each study design category prevents us from making definitive claims about species-specific SOC gains (Fig. S5, Table S1). Therefore, we did not find evidence that our pooled results may be biased by the type of animal used.

## 5. Conclusion

Our analysis finds little robust support for the hypothesis that regenerative grazing management can meaningfully sequester carbon in soil organic matter. Relatedly, there is little evidence that regenerative grazing could potentially offset an appreciable quantity of ruminant enteric fermentation methane or other greenhouse gas emissions. In order to test this hypothesis more rigorously, more case-controlled longitudinal studies are needed. At a minimum, past cross-sectional study sites would need only an additional soil measurement of similar quality, if previous grazing management methods have continued, to provide higher-quality evidence that SOC sequestration is occurring, and to attribute that SOC sequestration to regenerative grazing itself. Ideally, however, multiple more years of measurement should be conducted.

## Supporting information

Data S1 - Meta-Analysis Data

Data S2 - R script for plots and stats

## Author Contributions

**Docker Clark**: data curation (lead); formal analysis (supporting); investigation (equal); visualization (equal); writing - original draft preparation (equal). **Jasmine A. Dillon**: methodology (supporting); visualization (supporting); writing - review and editing (equal). **Matthew N. Hayek**: conceptualization (lead); data curation (supporting); formal analysis (lead); funding acquisition (lead); investigation (equal); methodology (lead); project administration (lead); supervision (lead); visualization (equal); writing - original draft preparation (equal); writing - review and editing (equal). **Shelby C. McClelland**: formal analysis (supporting); methodology (supporting); visualization (supporting); writing - review and editing (equal).

