## Supplementary information for "Regenerative Grazing as a Climate Change Mitigation Strategy: A Systematic Review"

**Contents:**

Box S1: Complete search terms …………………………………………………………….…. p.1

Figure S1: PRISMA Flow diagram …………………………………………………………..…… p. 2

Figure S2: Site-wise analysis of SOC flux ………………………………………………………..  p. 3

Figure S3: Normalized to 100 cm depth (study-wise) …………...………………………………. p. 4

Figure S4: Normalized to 30 cm depth (study-wise) ………………...………………………… p. 5

Figure S5: Grazed species analysis ……………………………...………………………… p. 6

Table S1: Species-wise summary statistics ………………………………….…………… p. 7

Figure S6: Non-continuous counterfactual (study-wise) ………………………………………… p. 8

Figure S7: Non-continuous counterfactual (site-wise) …………………………………………… p. 9

Supplementary Methods and Results …………………………….…………………………… p. 10

Sources ……………………………………………………………………………………..…. p. 11

*ALL=((Ruminants OR cattle OR cow OR beef OR milk OR sheep OR goats OR mutton OR wool) AND (pasture OR rangeland OR meadow OR prairie OR steppe) AND (“mob grazing” OR “savory method” OR “adaptive multi-paddock” OR “rotational grazing” OR “holistic management” OR “holistic planned grazing” OR “management-intensive grazing”) AND (“SOC” OR “soil carbon” OR “SOM” OR “soil organic matter” OR “soil organic material” OR “soil organic carbon” OR “soil organic content”))*

Box S1: Complete search criteria from WebofScience.com (search conducted Feb 5, 2025 ).

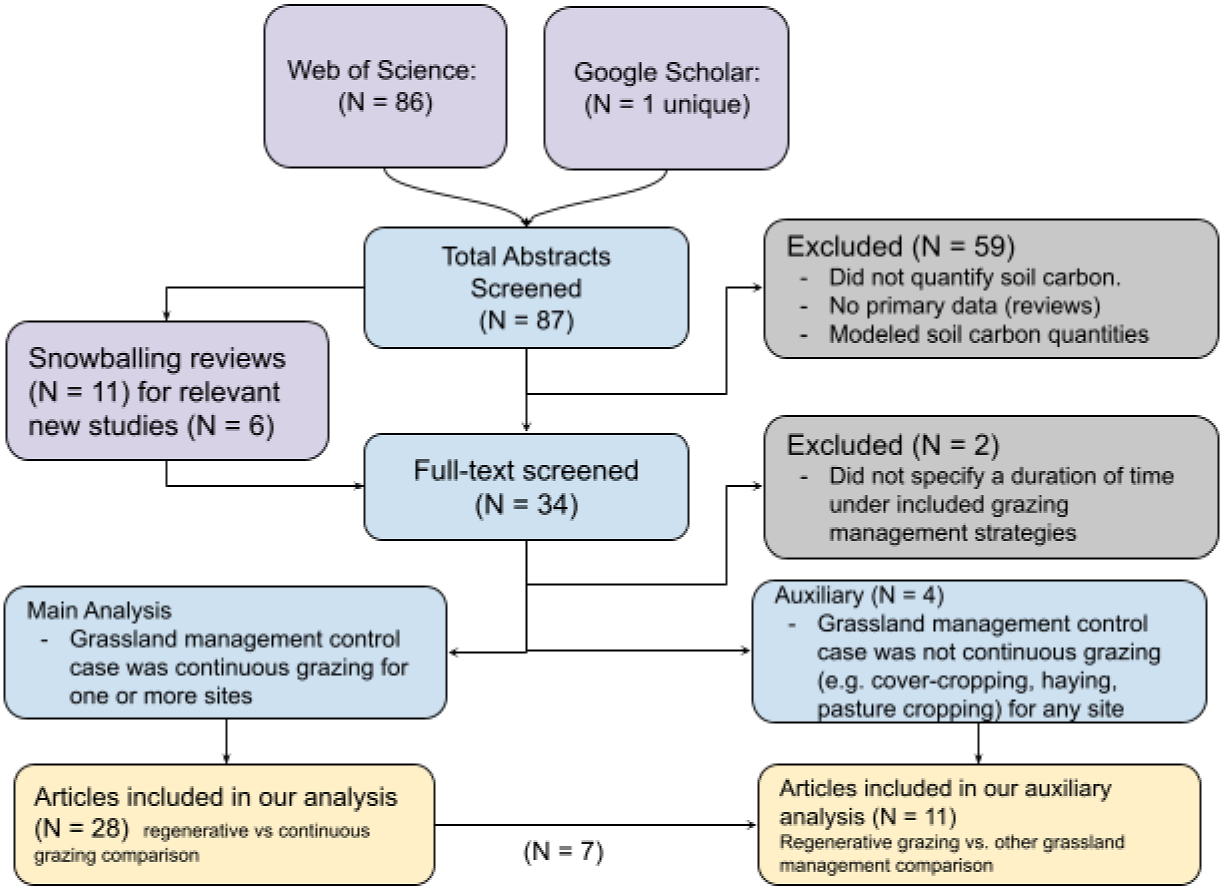


Fig S1. PRISMA flow diagram. Reasons for disqualification in gray boxes. Sources of studies in violet boxes.

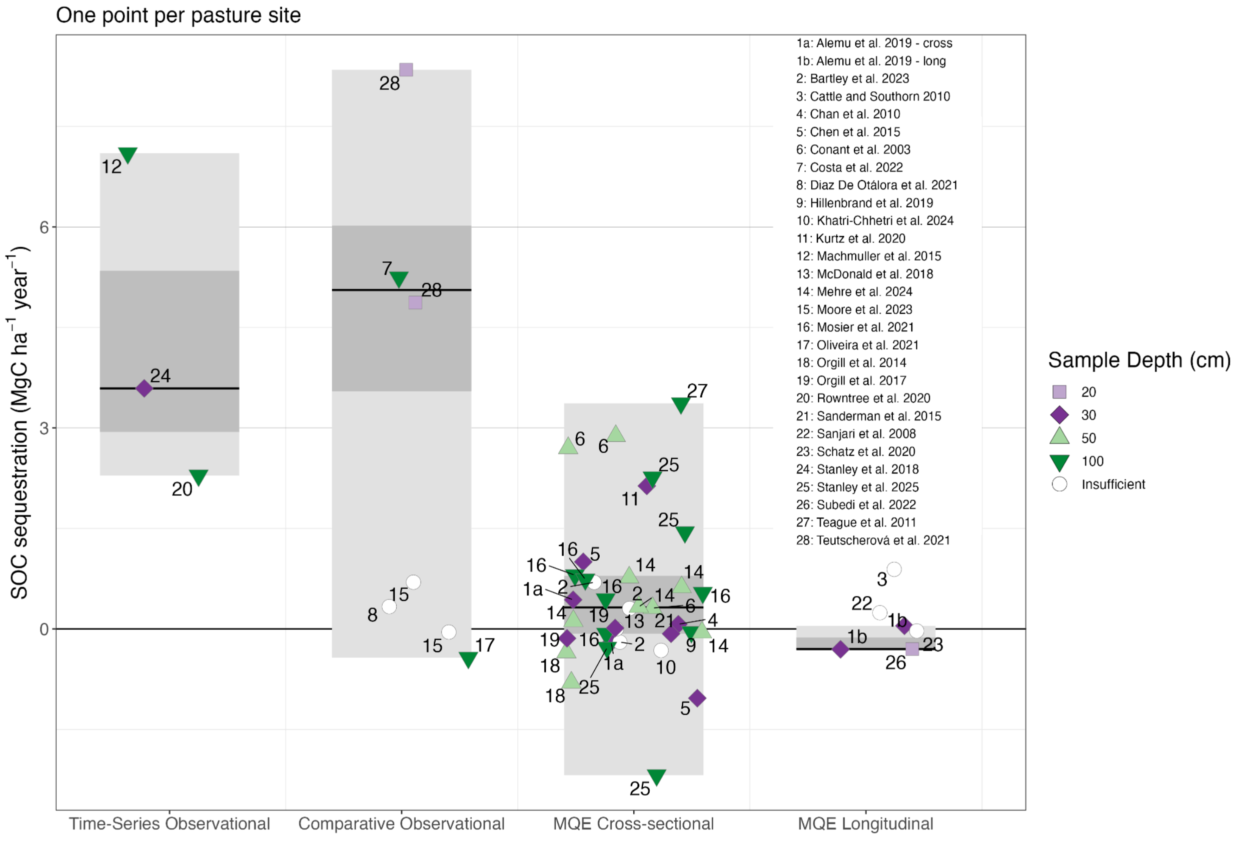


Fig. S2: SOC sequestration rates in each study-site pair (points) of each regenerative grazing study (numbers) and ranges. Colors represent the maximum depth at which SOC measurements were conducted. Studies with depth measurements of <20 cm or with only one depth stratum were classified as having insufficient depth data, for possible exclusion from analysis (white points). Light gray boxes represent full ranges excluding insufficient depth data. Dark gray boxes represent interquartile ranges, and black lines show medians for studies conducted with robust SOC depth data. The SOC sequestration flux was calculated as the difference in time or across chronosequence sample points in the time-series observational category, as the differences between paired plots in pasture sites (averaged over all sites in each study) divided by number of years of management differentiation in the MQE Cross-Sectional category, and as the difference-in-difference across paired plots and over time in the MQE Longitudinal category.

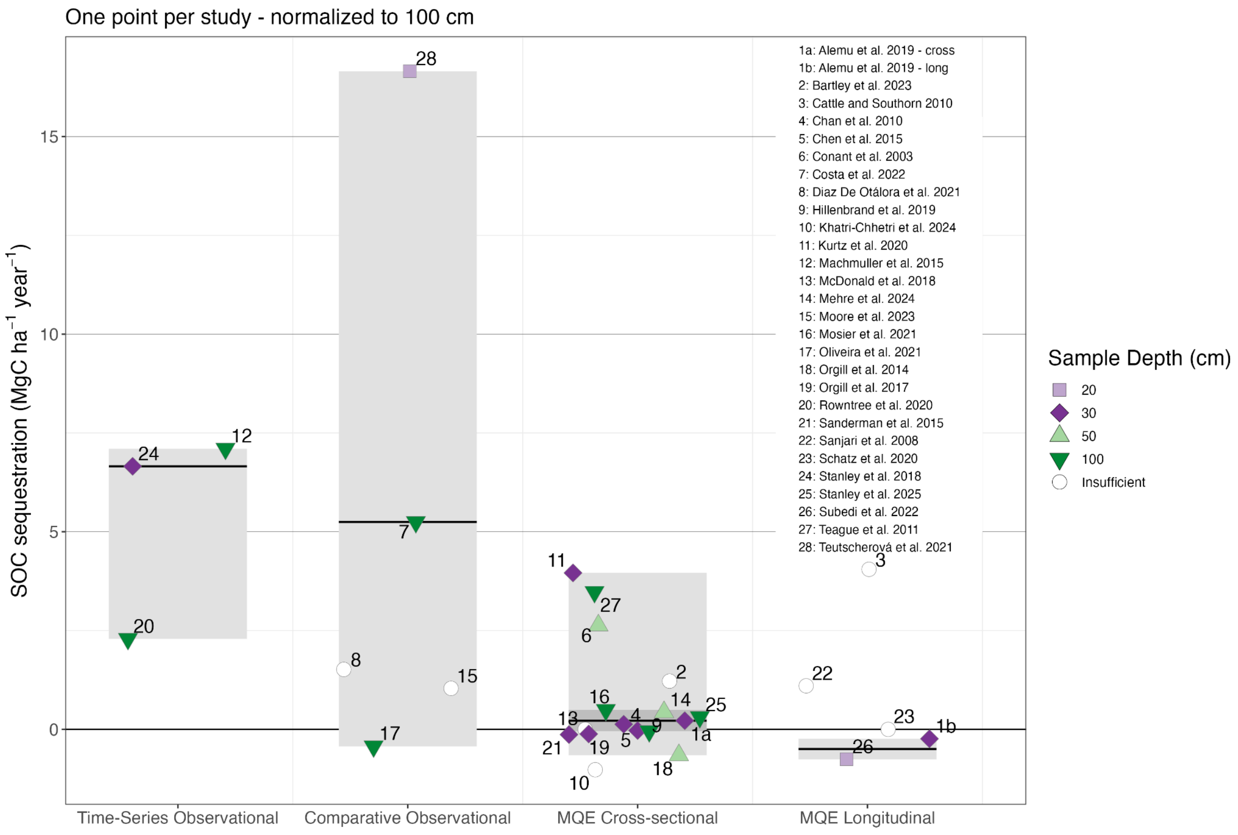


Fig. S3: Mean SOC sequestration rates in each regenerative grazing study (numbers) and ranges. SOC sequestration rates (points) are normalized to a common depth of 100cm using methods reported in McClelland et al. 2021. Colors represent the maximum depth at which SOC measurements were conducted. Studies with depth measurements of <20 cm or with only one depth stratum were classified as having insufficient depth data, for possible exclusion from analysis (white points). Light gray boxes represent full ranges excluding insufficient depth data. Dark gray boxes represent interquartile ranges, and black lines show medians for studies conducted with robust SOC depth data. The SOC sequestration flux was calculated as the as the difference in time or across chronosequence sample points in the time-series observational category, as the differences between paired plots in pasture sites (averaged over all sites in each study) divided by number of years of management differentiation in the MQE Cross-Sectional category, and as the difference-in-difference across paired plots and over time in the MQE Longitudinal category.

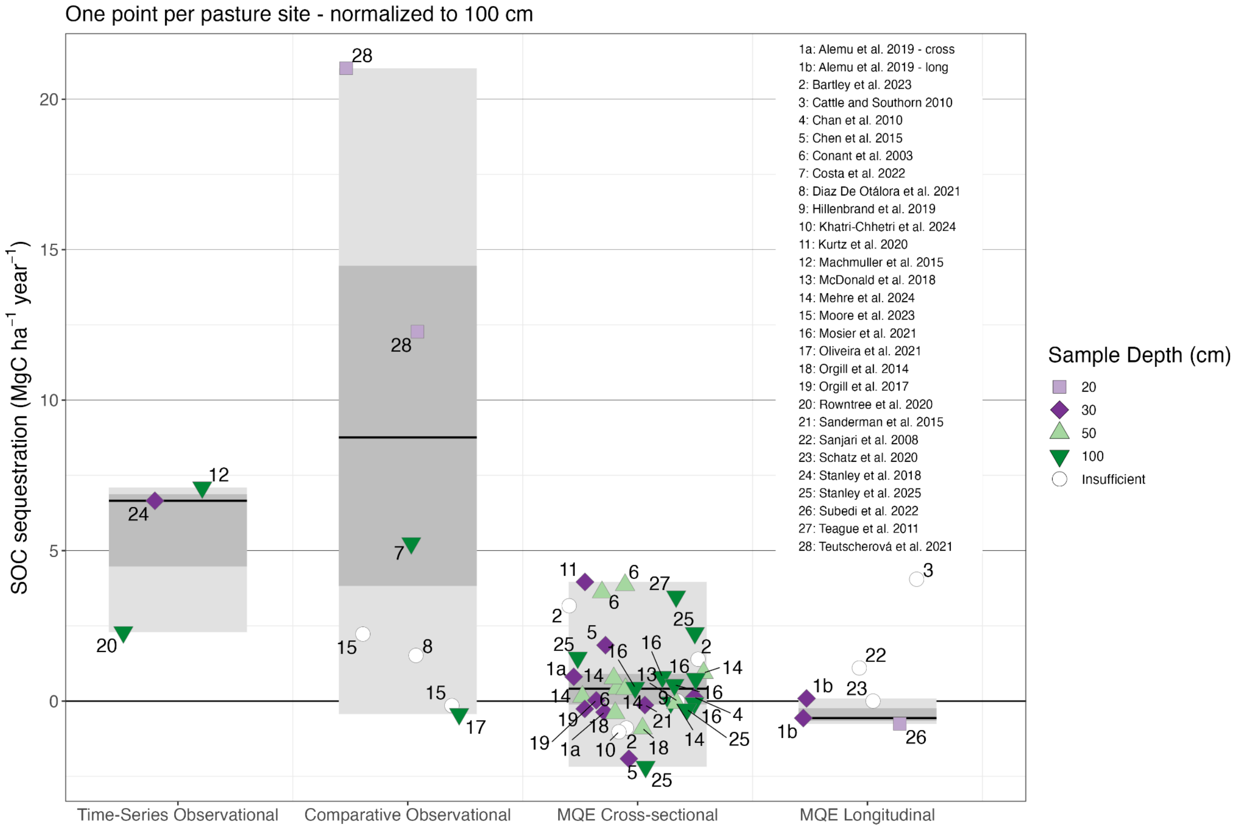


Fig. S4: SOC sequestration rates in each study site pair (points) of each regenerative grazing study (numbers) and ranges. SOC sequestration rates are normalized to a common depth of 100cm using methods reported in McClelland et al. 2021. Colors represent the maximum depth at which SOC measurements were conducted. Studies with depth measurements of <20 cm or with only one depth stratum were classified as having insufficient depth data, for possible exclusion from analysis (white points). Light gray boxes represent full ranges excluding insufficient depth data. Dark gray boxes represent interquartile ranges, and black lines show medians for studies conducted with robust SOC depth data. The SOC sequestration flux was calculated as the as the difference in time or across chronosequence sample points in the time-series observational category, as the differences between paired plots in pasture sites (averaged over all sites in each study) divided by number of years of management differentiation in the MQE Cross-Sectional category, and as the difference-in-difference across paired plots and over time in the MQE Longitudinal category.

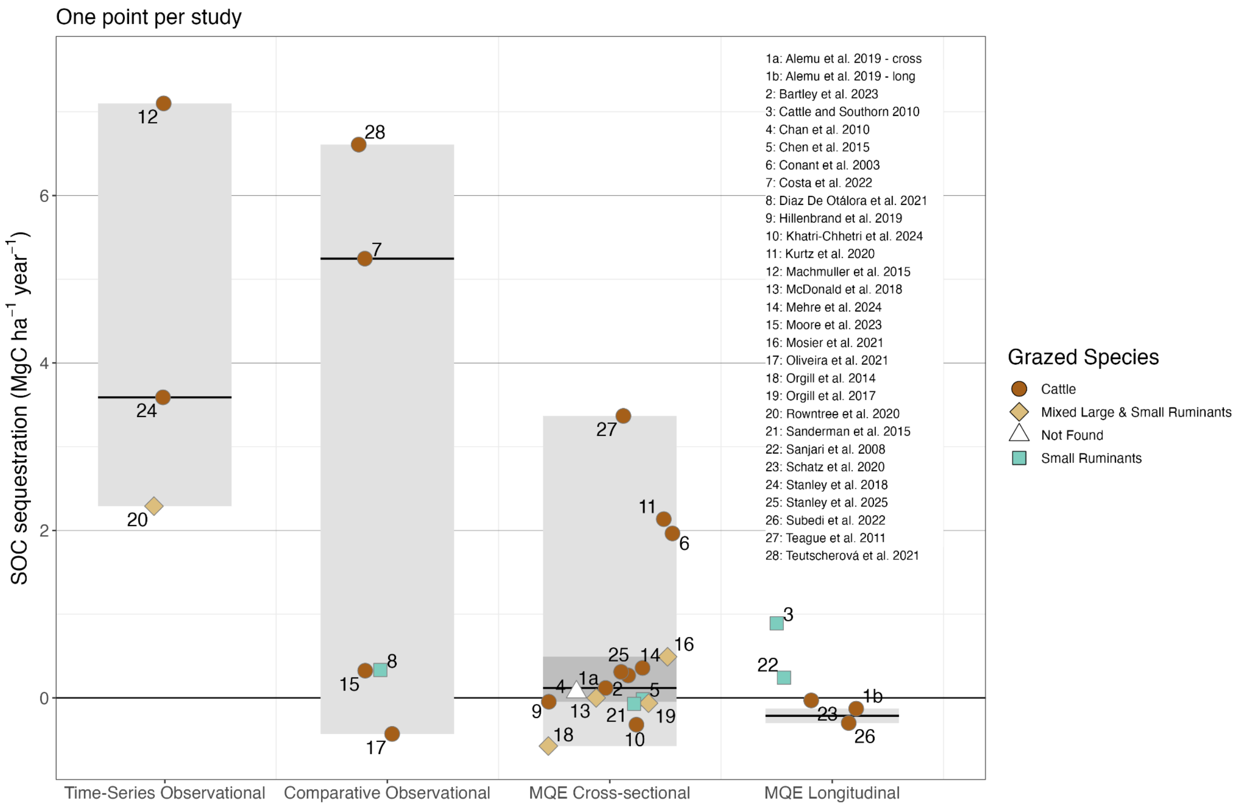


Fig. S5: Mean SOC sequestration rates in each regenerative grazing study (numbers) and ranges. Colors and point shapes represent the domestic species grazed. Studies with depth measurements of <20 cm or with only one depth stratum were classified as having insufficient depth data, for possible exclusion from analysis. Light gray boxes represent full ranges excluding insufficient depth data. Dark gray boxes represent interquartile ranges, and black lines show medians for studies conducted with robust SOC depth data. One study (Chan et al. 2010) did not report the species grazing their pastures and was placed into the “Not Found” category (white point). The SOC sequestration flux was calculated as the as the difference in time or across chronosequence sample points in the observational category, as the differences between paired plots in pasture sites (averaged over all sites in each study) divided by number of years of management differentiation in the Cross-Sectional category, and as the difference-in-difference across paired plots and over time in the Longitudinal category.

| Species | Studies | Mean | Median | Range |
|---|---|---|---|---|
| **Cattle** | 16 | 1.61 | 0.32 | -0.43 - 7.1 |
| **Mixed Large & Small Ruminants** | 5 | 0.43 | 0 | -0.58 - 2.29 |
| **Small Ruminants** | 5 | 0.28 | 0.24 | -0.07 - 0.89 |
| **Not Found** | 1 | 0.07 | 0.07 | 0.07 - 0.07 |

Table S1: Summary statistics and ranges for SOC sequestration (positive numbers = sequestration) categorized by grazed species. Small ruminants are either sheep, goats, or a combination thereof while mixed large and small ruminants encompasses any combination of cattle with sheep or goats. One study (Chan et al. 2010) did not contain enough information on species to confidently categorize.

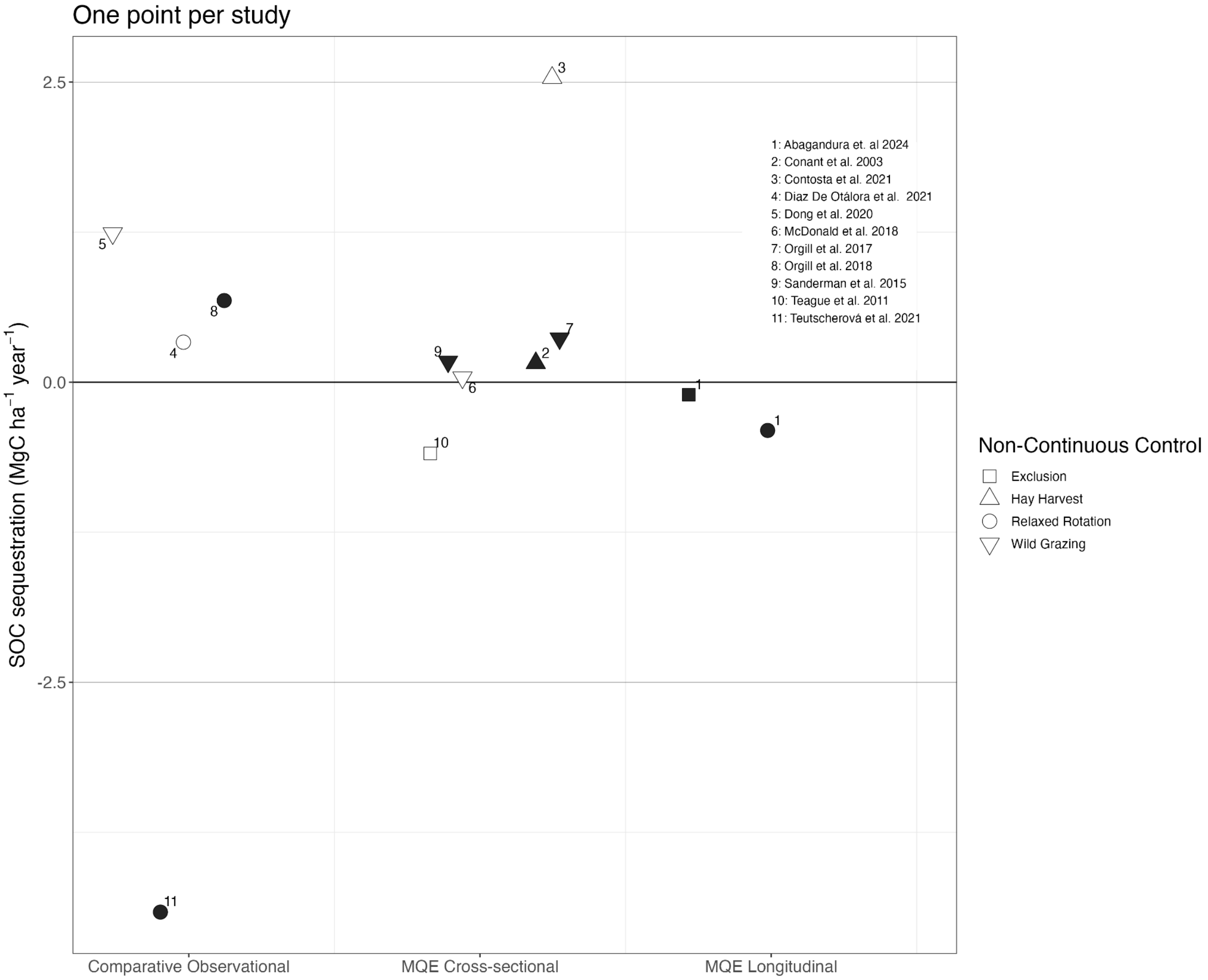


Fig. S6: Mean SOC sequestration in each regenerative grazing study (numbers) that used a control site which was not strictly continuous grazing. Shapes represent the type of non-continuous control used for comparison to regeneratively grazed sites. Studies with depth measurements of <20 cm or with only one depth stratum were classified as having insufficient depth data, for possible exclusion from analysis (white points).

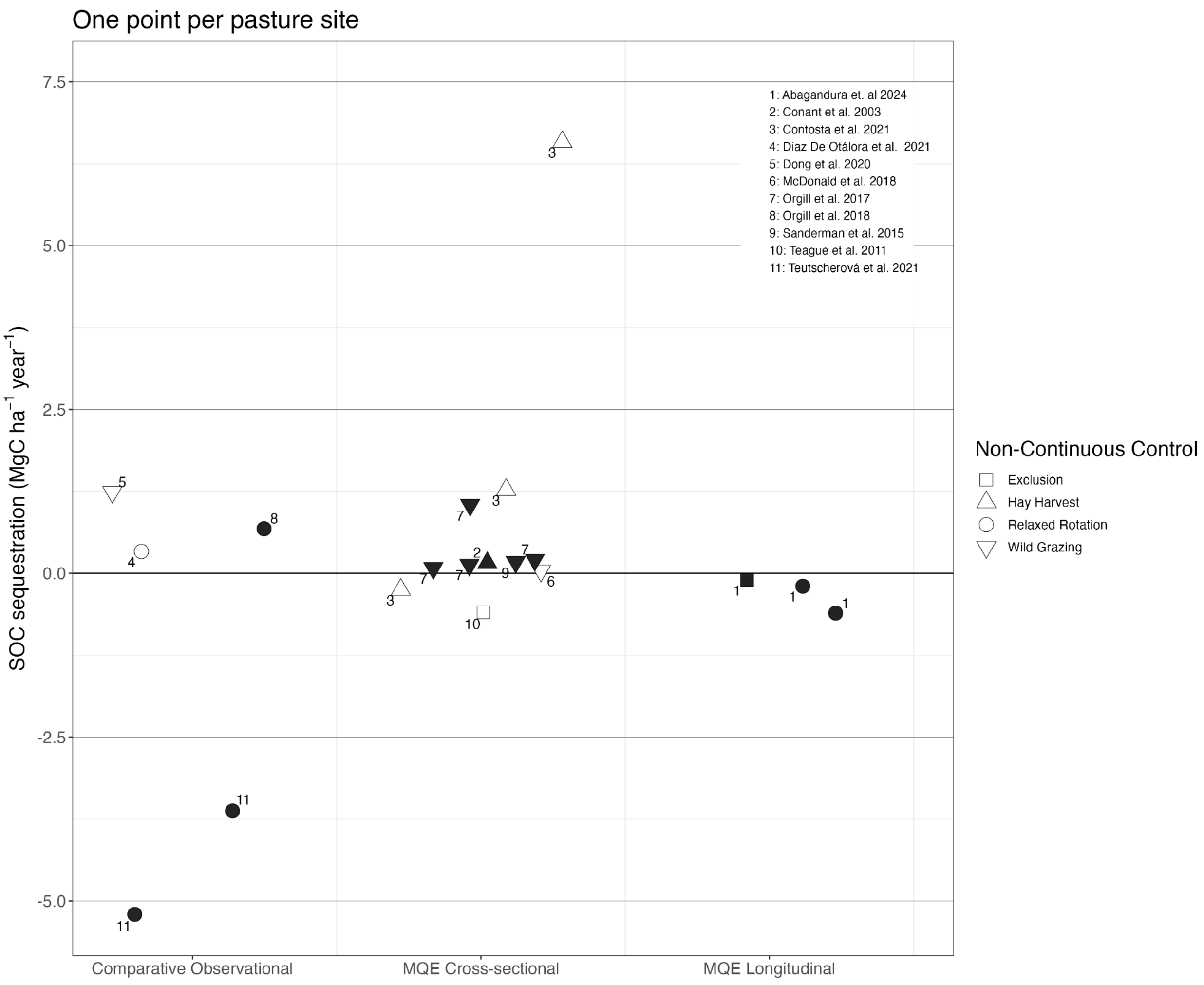


Fig. S7: SOC sequestration in each regenerative grazing study (numbers) that used a control site which was not strictly continuous grazing. Points represent regenerative-non-conventional study site pairs. Shapes represent the type non-continuous control used for comparison to regeneratively grazed sites. Studies with depth measurements of <20 cm or with only one depth stratum were classified as having insufficient depth data, for possible exclusion from analysis (white points).

### 1. Supplementary methods and results

#### 1.1 Differences between study-calculated SOC fluxes and our methods.

There are cases wherein parent studies report SOC flux in a way that does not provide direct case-control between regeneratively grazed and conventionally grazed sites (i.e. Oliveira et al. 2021). This study compared SOC under three management strategies: continuous low-stocking grazing (CLS), rotational high-stocking grazing (RHS), and neighboring native Atlantic Forest (FOR). The authors calculated carbon accumulation rates (CAR) by subtracting the FOR carbon stock (assumed at equilibrium) from CLS and RHS stocks, then dividing by management duration (28 and 18 years, respectively), yielding CARs of 0.45 and 0.02 Mg C $ha^{-1}$ $yr^{-1}$. Our analysis uses the difference between these rates (-0.43 Mg C $ha^{-1}$ $yr^{-1}$).

However, this approach presents methodological limitations. First, it relies on indirect comparison using the forest reference rather than temporal SOC measurements within each grazing system. Second, the assumption that forest soils represent equilibrium conditions is questionable, given that these sites likely experienced historical land-use changes from forest to pasture, with associated SOC impacts.

For direct comparison, we recalculated CAR by subtracting CLS from RHS carbon stocks and dividing by average management duration (23 years), yielding -0.53 Mg C $ha^{-1}$ $yr^{-1}$. While this adjustment does not materially affect our results, it eliminates dependence on the forest reference and provides a more robust treatment-control comparison.

#### 1.2 Livestock species as a predictor variable

To evaluate the potential for farmed livestock species as a confounding variable (Fig. S5), we calculated summary statistics for our dataset separating categories by grazed species (Table S1). A Mann-Whitney U test was used to evaluate the significance of SOC sequestration on cattle-grazed vs all other ruminant grazed categories (Mixed Large and Small + Small Ruminants). This returned a p-value of 0.0879, which is not significant at the 95% confidence level. We performed this test on only studies within the MQE cross-sectional category as this provided the largest and most even sample size of cattle-grazing (N = 9) and non-cattle ruminant grazing (N = 6) studies.

**SI Sources:**


Abagandura, G. O., Mamo, M., Schacht, W. H., Shropshire, A., & Volesky, J. D. (2024). Soil carbon and nitrogen after eight years of rotational grazing in the Nebraska Sandhills meadows. *Geoderma*, *442*, 116776. https://doi.org/10.1016/j.geoderma.2024.116776

Abdalla, M., Hastings, A., Cheng, K., Yue, Q., Chadwick, D., Espenberg, M., Truu, J., Rees, R. M., & Smith, P. (2019). A critical review of the impacts of cover crops on nitrogen leaching, net greenhouse gas balance and crop productivity. *Global Change Biology*, *25*(8), 2530–2543. https://doi.org/10.1111/gcb.14644

Alexander, P., Brown, C., Arneth, A., Finnigan, J., Moran, D., & Rounsevell, M. D. A. (2017). Losses, inefficiencies and waste in the global food system. *Agricultural Systems*, *153*, 190–200. https://doi.org/10.1016/j.agsy.2017.01.014

Beillouin, D., Corbeels, M., Demenois, J., Berre, D., Boyer, A., Fallot, A., Feder, F., & Cardinael, R. (2023). A global meta-analysis of soil organic carbon in the Anthropocene. *Nature Communications*, *14*(1), 3700. https://doi.org/10.1038/s41467-023-39338-z

Byrnes, R. C., Eastburn, D. J., Tate, K. W., & Roche, L. M. (2018). A Global Meta‐Analysis of Grazing Impacts on Soil Health Indicators. *Journal of Environmental Quality*, *47*(4), 758–765. https://doi.org/10.2134/jeq2017.08.0313

Conant, Richard T., Carlos E. P. Cerri, Brooke B. Osborne, and Keith Paustian. “Grassland Management Impacts on Soil Carbon Stocks: A New Synthesis.” Ecological Applications 27, no. 2 (2017): 662–68. https://doi.org/10.1002/eap.1473.

Contosta, A. R., Arndt, K. A., Campbell, E. E., Stuart Grandy, A., Perry, A., & Varner, R. K. (2021). Management intensive grazing on New England dairy farms enhances soil nitrogen stocks and elevates soil nitrous oxide emissions without increasing soil carbon. *Agriculture, Ecosystems & Environment*, *317*, 107471. https://doi.org/10.1016/j.agee.2021.107471

Gosnell, H., Charnley, S., & Stanley, P. (2020). Climate change mitigation as a co-benefit of regenerative ranching: insights from Australia and the United States. *Interface Focus*, *10*(5), 20200027. https://doi.org/10.1098/rsfs.2020.0027

Henry, B., Allen, D., Badgery, W., Bray, S., Carter, J., Dalal, R. C., Hall, W., Harrison, M. T., McDonald, S. E., & McMillan, H. (2024). Soil carbon sequestration in rangelands: a critical review of the impacts of major management strategies. *The Rangeland Journal*, *46*(3). https://doi.org/10.1071/RJ24005

Jobbágy, E. G., & Jackson, R. B. (2000). THE VERTICAL DISTRIBUTION OF SOIL ORGANIC CARBON AND ITS RELATION TO CLIMATE AND VEGETATION. *Ecological Applications*, *10*(2), 423–436. https://doi.org/10.1890/1051-0761(2000)010%5B0423:tvdoso%5D2.0.co;2

McClelland, S. C., Paustian, K., & Schipanski, M. E. (2021). Management of cover crops in temperate climates influences soil organic carbon stocks: a meta-analysis. *Ecological Applications*, *31*(3), e02278. https://doi.org/10.1002/eap.2278

McDonald, S. E., Badgery, W., Clarendon, S., Orgill, S., Sinclair, K., Meyer, R., Butchart, D. B., Eckard, R., Rowlings, D., Grace, P., Doran-Browne, N., Harden, S., Macdonald, A., Wellington, M., Pachas, A. N. A., Eisner, R., Amidy, M., & Harrison, M. T. (2023). Grazing management for soil carbon in Australia: A review. *Journal of Environmental Management*, *347*, 119146. https://doi.org/10.1016/j.jenvman.2023.119146

Oliveira, P. P. A., Rodrigues, P. H. M., Praes, M. F. F. M., Pedroso, A. F., Oliveira, B. A., Sperança, M. A., Bosi, C., & Fernandes, F. A. (2021). Soil carbon dynamics in Brazilian Atlantic forest converted into pasture-based dairy production systems. *Agronomy Journal*, *113*(2), 1136–1149. https://doi.org/10.1002/agj2.20578

Orgill, S. E., Condon, J. R., Conyers, M. K., Morris, S. G., Alcock, D. J., Murphy, B. W., & Greene, R. S. B. (2018). Removing Grazing Pressure from a Native Pasture Decreases Soil Organic Carbon in Southern New South Wales, Australia. *Land Degradation & Development*, *29*(2), 274–283. https://doi.org/10.1002/ldr.2560

Stanley, P. L., Wilson, C., Patterson, E., Machmuller, M. B., & Cotrufo, M. F. (2024). Ruminating on soil carbon: Applying current understanding to inform grazing management. *Global Change Biology*, *30*(3), e17223. https://doi.org/10.1111/gcb.17223

Wang, J., Li, Y., Bork, E. W., Richter, G. M., Chen, C., Hussain Shah, S. H., & Mezbahuddin, S. (2021). Effects of grazing management on spatio-temporal heterogeneity of soil carbon and greenhouse gas emissions of grasslands and rangelands: Monitoring, assessment and scaling-up. *Journal of Cleaner Production*, *288*, 125737. https://doi.org/10.1016/j.jclepro.2020.125737

Wang, X., McConkey, B. G., VandenBygaart, A. J., Fan, J., Iwaasa, A., & Schellenberg, M. (2016). Grazing improves C and N cycling in the Northern Great Plains: a meta-analysis. *Scientific Reports*, *6*(1), 33190. https://doi.org/10.1038/srep33190

Zhou, G., Zhou, X., He, Y., Shao, J., Hu, Z., Liu, R., Zhou, H., & Hosseinibai, S. (2017). Grazing intensity significantly affects belowground carbon and nitrogen cycling in grassland ecosystems: a meta‐analysis. *Global Change Biology*, *23*(3), 1167–1179. https://doi.org/10.1111/gcb.13431
