## Supplementary material for "Regenerative Grazing as a Climate Change Mitigation Strategy: A Systematic Review": Data S2 - R script for plots and stats: Data_S2-SOC_flux_figs.R

```
# Created by Docker B. Clark and Matthew N. Hayek
# Saved and uploaded 10/30/2025


library(dplyr)
library(ggplot2)
library(matrixStats)
#install.packages("readxl")

#========================== Regen Ag SOC data review ================================# 
# setwd("/Users/dockerclark/Desktop/Work/RegenAg NYU/Data/regengrazing") #Docker
# setwd("~/Dropbox/Research/Modeling/Regen_ag/") #Matthew

# Packages
require(tidyverse)
require(ggrepel)
# require(rgb)
require(readxl)

# Data
Sequest <- read_excel("Regenerative Grazing SOC Meta-Analysis.xlsx", 
                      sheet = "Continuous grazing counterfact")


# Rename Norm.diff.flux to Norm.diff.flux.30 to specify it's 30 cm normalization
Sequest <- Sequest %>%
    rename(Norm.diff.flux.30 = Norm.diff.flux)

#### Data Tidying & Manipulation ####
# run these all once

# Add Citation column to display author & year at once
Sequest$Citation <- paste(Sequest$Author, Sequest$Year, sep = " ")
Sequest$Citation <- factor(Sequest$Citation)
# Remove the archived studies
Sequest <- Sequest %>%
  filter(Archive == FALSE)

# rename all continents besides NAm, SAm, and Aus to "Other" and relevel
Sequest <- Sequest %>%
    mutate(
        Region = Region %>%
               #  collapse all other continents into "Other"
               forcats::fct_other( keep = c("North America", "South America", "Australia"),
                   other_level = "Other") %>%
    forcats::fct_relevel("North America", "South America", "Australia", "Other")
    )

# Pull out ONLY difference fluxes
Fluxes <- Sequest %>%
  filter(Difference.flux.tf == TRUE)


# Reorder levels of Cat to change column order
Fluxes$Cat <- factor(Fluxes$Cat, 
                     levels = c("Time-Series Observational", "Comparative Observational", 
                                "MQE Cross-Sectional", "MQE Longitudinal"))


# specify visualized depth levels
depth_levels <- c(5, 10, 20, 30, 50, 100)

# sum all depth slices and round to nearest depth level specified above
Fluxes <- Fluxes %>%
  mutate(Sample.Depth.cm = select(., starts_with("Slice")) %>%
           rowSums(na.rm = TRUE)) %>%
  mutate(Sample.Depth.cm = map_dbl(Sample.Depth.cm,
                                   ~depth_levels[which.min(abs(depth_levels - .x))]))

# create a column that counts depth slices
Fluxes <- Fluxes %>%
  mutate(Slices = str_count(All_depths.cm, ",") + 1)

# Create a jittered x-value column to have predictable jitter
set.seed(07072025) # date of coding setting a seed for reproducability
Fluxes$jittered_x <- jitter(as.numeric(Fluxes$Cat), amount = 0.3)

# Create a binary variable for robust depth info
Fluxes$depth.tf <- ifelse(Fluxes$Sample.Depth.cm > 10 & 
                            Fluxes$Slices > 1, TRUE, FALSE)

# Allow for custom coloring later
Fluxes$depth_factor <- factor(
  ifelse(Fluxes$depth.tf == FALSE, "Insufficient", as.character(Fluxes$Sample.Depth.cm)),
  levels = c("20", "30", "50", "100", "Insufficient"))


# getting unique citation
unique_citations <- Fluxes %>%
  group_by(Citation) %>%
  summarise(Citation = first(Citation)) # get all unique citations

# give each unique citation a number 
unique_citations$Label <- c("1a", "1b", 
                            as.character(2:(length(unique_citations$Citation)-1)))


# left-join by shared "Citation". result should work for site and study wise plots
Fluxes <- Fluxes %>% 
  left_join(unique_citations, by = "Citation")

Fluxes <- Fluxes %>% # adding a text column for the legend
  mutate(citation_display = paste0(Citation, " - ", Label)) %>%
  # Create a sorting key based on Label 
  mutate(label_numeric = as.numeric(gsub("[a-z]", "", Label)),
         
         label_letter = gsub("[0-9]", "", Label)) %>%
  
  arrange(label_numeric, label_letter) %>%
  # levels are arranged first by number then letter
  
  mutate(citation_display = factor(citation_display, levels = unique(citation_display))) %>%

  # remove temporary sorting columns 
  select(-label_numeric, -label_letter)


# Create a df of study-wise difference fluxes
Fluxes.study <- Fluxes %>%
  group_by(Citation) %>%
  # using first() to retain any necessary columns from original df
  summarize(Author = first(Author), Year = first(Year), Cat = first(Cat), 
            All_depths.cm = first(All_depths.cm), Sample.Depth.cm = first(Sample.Depth.cm),
            Slices = first(Slices), jittered_x = first(jittered_x), depth.tf = first(depth.tf),
            depth_factor = first(depth_factor), Significant.tf = first(Significant.tf),
            Region = first(Region), Significance_tested.tf = first(Significance_tested.tf),
            Species = first(Species), Flux.Discussion = first(Flux.Discussion),
            Label = first(Label), citation_display = first(citation_display),
            OG.unit = first(OG.unit), causal = first(Causal.Language), 
            # mean flux for each study
            Flux.Mghayr = mean(Flux.Mghayr),
            Norm.diff.flux.30 = mean(Norm.diff.flux.30),
            Norm.diff.flux.100 = mean(Norm.diff.flux.100))


# Change this for different output
# select "studies" or "sites"
data_type <- "studies"

if (data_type == "studies"){
    flux.df <- Fluxes.study ; print("data type set to 'studies'")
    data_title <- "One point per study"
} else if (data_type=="sites"){
    flux.df <- Fluxes ; print("data type set to 'sites'")
    data_title <- "One point per pasture site"
} else { 
    flux.df <- NULL ; print("invalid selection for data type")
    data_title <- NULL
}

# select "original", "30", or "100" cm
normalization <- "original"

if (normalization == "original"){
    flux.df <- flux.df %>% 
        mutate(SOC_flux = Flux.Mghayr)
     normal_title = NULL
     print("no normalization")
} else if (normalization == "30"){
    flux.df <- flux.df %>% 
        mutate(SOC_flux = Norm.diff.flux.30)
    normal_title = " - normalized to 30 cm"
    print(" normalization")
} else if (normalization == "100"){
    flux.df <- flux.df %>% 
        mutate(SOC_flux = Norm.diff.flux.100)
    normal_title = " - normalized to 100 cm"
    print("normalization = 100 cm")
} else { 
    flux.df <- flux.df %>% 
        mutate(SOC_flux =NULL)
    normal_title = NULL
    print("invalid selection for normalization ")
}


# an option to re-roll the jitter
if (data_type == "studies" & (normalization == "original" | normalization == "30")) {

  set.seed(072025) 
  #overwrite jittered x with new seed
  flux.df$jittered_x <- jitter(as.numeric(flux.df$Cat), amount = 0.3) 
  print("rerolled jitter position to preset seed")
  
} else if (data_type == "studies" & normalization == "100") {
  #Studies norm100
  set.seed(987654) #StudiesNorm100
  flux.df$jittered_x <- jitter(as.numeric(flux.df$Cat), amount = 0.3)
  print("rerolled jitter position to preset seed")
  
} else if (data_type == "sites" & normalization == "original") {
  #sites norm = original
  set.seed(07072025) 
  flux.df$jittered_x <- jitter(as.numeric(flux.df$Cat), amount = 0.3) 
  print("rerolled jitter position to preset seed")
  
} else if (data_type == "sites" & normalization == "30") {
  #sites norm = 30
  set.seed(5) #Sitesnorm30
  flux.df$jittered_x <- jitter(as.numeric(flux.df$Cat), amount = 0.3) 
  print("rerolled jitter position to preset seed")
  
} else if (data_type == "sites" & normalization == "100") {
  #sites norm = 100
  set.seed(1998) #sitesnorm100
  flux.df$jittered_x <- jitter(as.numeric(flux.df$Cat), amount = 0.3)
  print("rerolled jitter position to preset seed")
}

depth_colors <- c("#c2a5cf", "#7b3294",  "#a6dba0", "#008837","white") #from colorbrewer2

#filter out excluded depth data for plots and stats
excluded <- flux.df %>%
  filter(depth.tf == TRUE)

# creating a df for whisker caps
minmax <- excluded %>%
  group_by(Cat) %>%
  summarise(min = as.numeric(min(SOC_flux)), 
         max = as.numeric(max(SOC_flux)))
minmax <- cbind(minmax, c(1, 2, 3, 4))
colnames(minmax) <- c("Cat", "min", "max", "x")

citation_legend <- flux.df %>%
  select(Label, Citation) %>%
  distinct() %>%
  arrange(as.numeric(gsub("[a-z]", "", Label))) %>%  # Sort numerically
  mutate(legend_text = paste0(Label, ": ", Citation)) %>%
  pull(legend_text) %>%
  paste(collapse = "\n")

boxcolor <- "gray75"
boxlines <- 1.5
boxwidth <- 0.50

excl_stats <- excluded %>%
    mutate(catNum = as.numeric(Cat)) %>%
    group_by(catNum) %>%
    summarise(
        medn = median(SOC_flux, na.rm = TRUE),
        min = min(SOC_flux, na.rm = TRUE),
        max = max(SOC_flux, na.rm = TRUE),
        q25 = quantile(SOC_flux, 0.25, na.rm = TRUE),
        q75 = quantile(SOC_flux, 0.75, na.rm = TRUE)) %>%
    mutate(q25 = q25, q75 =  q75)


####--------------------------Plots with DEPTH colors---------------------------#### 
flux.depth.gg <- ggplot(flux.df) +

  geom_abline(mapping = aes(intercept = 0, slope = 0))

if (data_type == "studies") {
  flux.depth.gg <- flux.depth.gg +
    
    #plots for cats with 3 or more points
      geom_rect(data = excl_stats,
                aes(xmin = catNum - 0.3, xmax = catNum + 0.3,
                    ymin = min, ymax = max),
                fill = "gray90",
                color = NA) +  
      geom_rect(data = filter(excl_stats, catNum == 3),
                aes(xmin = catNum - 0.3, xmax = catNum + 0.3,
                    ymin = q25, ymax = q75),
                fill = "gray75",
                color = NA) +

      geom_segment(data = excl_stats,
                   aes(x = catNum - 0.3, xend = catNum + 0.3,
                       y = medn, yend = medn),
                   color = "black",
                   linewidth = 0.75) 
      
      
      
      
   #  geom_col(data = subset(excluded), # Only draw bars for the non-insuff studies
   #             aes(x = Cat, y = SOC_flux),
   #             position = "identity", width = boxwidth, fill = "gray90") + 
   #  geom_boxplot(data = filter(excluded, Cat != "MQE Longitudinal"), 
   #               mapping = aes(x = Cat, y = SOC_flux), width = boxwidth, 
   #               coef = 0, alpha = 0, 
   #               linewidth = boxlines,color=NULL, fill = "gray75")  

      
      # geom_linerange(data = filter(minmax, Cat == "MQE Longitudinal"),
      #              mapping = aes(x = Cat, ymin = min, ymax = max), 
      #              linewidth = boxlines, color = boxcolor) +
    # annotate("segment", 
    #          x = 4-boxwidth/2, 
    #          xend = 4+boxwidth/2,
    #          y = (minmax[4,2]+minmax[4,3])/2,
    #          yend = (minmax[4,2]+minmax[4,3])/2,
    #          linewidth = boxlines, color = boxcolor)
  
} else if (data_type == "sites") {
  flux.depth.gg <- flux.depth.gg +
    
    geom_rect(data = excl_stats,
              aes(xmin = catNum - 0.3, xmax = catNum + 0.3,
                  ymin = min, ymax = max),
              fill = "gray90",
              color = NA) +  
    geom_rect(data = excl_stats,
              aes(xmin = catNum - 0.3, xmax = catNum + 0.3,
                  ymin = q25, ymax = q75),
              fill = "gray75",
              color = NA) +
    
    geom_segment(data = excl_stats,
                 aes(x = catNum - 0.3, xend = catNum + 0.3,
                     y = medn, yend = medn),
                 color = "black",
                 linewidth = 0.75) 
}

#continue building the plot (this applies to both conditions)
flux.depth.gg <- flux.depth.gg +
  
  # #add the whisker caps on  boxplots
  # annotate("segment", y = minmax$min, yend = minmax$min, #bottom whiskers
  #          x = minmax$x-(boxwidth/2), xend = minmax$x+(boxwidth/2), 
  #          linewidth = boxlines, color = boxcolor) +
  # annotate("segment", y = minmax$max, yend = minmax$max, #top whiskers
  #          x = minmax$x-(boxwidth/2), xend = minmax$x+(boxwidth/2), 
  #          linewidth = boxlines, color = boxcolor) +
  
  #Points of different shape (and fill) to represent replicates (studies or sites)
  geom_point(aes(x = jittered_x, y = SOC_flux, 
                 fill = depth_factor, shape = depth_factor, color = depth.tf), 
             size = 5, stroke = 0.2) + 
  
  #specify color from above
  scale_fill_manual(values = depth_colors, name = "Sample Depth (cm)") +
  

  scale_color_manual(values = c(T = "white", F = "black")) +
  
  #specify manual shapes
  scale_shape_manual(values = c("20" = 22, "30" = 23, "50" = 24,
                                "100" = 25, "Insufficient" = 21),
                     name = "Sample Depth (cm)") +
  
  #add point labels
  geom_text_repel(aes(x = jittered_x, y = SOC_flux,
                      label = as.character(Label)),
                  size = 5, segment.size = 0.3,
                  max.overlaps = Inf) + 
  
  # Themes
  theme_bw() +
  theme(aspect.ratio = 0.8, text = element_text(size = 16),
        panel.grid.major.x = element_blank(),
        panel.grid.major.y = element_line(linewidth = .1, color = "black"),
        legend.text =  element_text(size = 10)) +
  
  #X axis labels
  scale_x_continuous(breaks = c(1, 2, 3, 4),
                   labels = c("Time-Series Observational", "Comparative Observational",
                              "MQE Cross-sectional", "MQE Longitudinal"))

#add conditional annotation based on data_type and normalization
if (data_type == "studies" & normalization == "original") {
  flux.depth.gg <- flux.depth.gg +
    #study wise, no norm label placement
    annotate("rect", 
             xmin = 3.6, ymin = 1.4, 
             xmax = 4.5, ymax = 7.51, 
             fill = "white") +
    #text
    annotate("text", 
             x = 3.7, y = 1.6, 
             label = citation_legend,
             hjust = 0, vjust = 0,
             size = 3.25,
             color = "black")
  
} else if (data_type == "studies" & normalization == "30") {
  flux.depth.gg <- flux.depth.gg +
    #study wise, norm 30 label placement
    annotate("rect", 
             xmin = 3.6, ymin = 1.4, 
             xmax = 4.5, ymax = 7.51, 
             fill = "white") +
    #text
    annotate("text", 
             x = 3.7, y = 1.6, 
             label = citation_legend,
             hjust = 0, vjust = 0,
             size = 3.25,
             color = "black")
  
} else if (data_type == "studies" & normalization == "100") {
  flux.depth.gg <- flux.depth.gg +
    #studies normalized to 100
    annotate("rect", 
             xmin = 3.6, ymin = 4.99, 
             xmax = 4.5, ymax = 15.5, 
             fill = "white") +
    annotate("text", 
             x = 3.7, y = 4.5, 
             label = citation_legend,
             hjust = 0, vjust = 0,
             size = 3.25,
             color = "black")
  
} else if (data_type == "sites" & normalization == "original") {
  flux.depth.gg <- flux.depth.gg +
    #sites and norm original
    annotate("rect", 
             xmin = 3.6, ymin = 1.4, 
             xmax = 4.5, ymax = 7.65, 
             fill = "white") +
    annotate("text", 
             x = 3.7, y = 1.26, 
             label = citation_legend,
             hjust = 0, vjust = 0,
             size = 3.25,
             color = "black")
  
} else if (data_type == "sites" & normalization == "30") {
  flux.depth.gg <- flux.depth.gg +
    #sites and norm = 30
    annotate("rect", 
             xmin = 3.6, ymin = 2.4, 
             xmax = 4.5, ymax = 11.3, 
             fill = "white") +
    annotate("text", 
             x = 3.7, y = 2.5, 
             label = citation_legend,
             hjust = 0, vjust = 0,
             size = 3.25,
             color = "black")
  
} else if (data_type == "sites" & normalization == "100") {
  flux.depth.gg <- flux.depth.gg +
    #sites and norm = 100
    annotate("rect", 
             xmin = 3.6, ymin = 4.9, 
             xmax = 4.5, ymax = 20.1, 
             fill = "white") +
    annotate("text", 
             x = 3.7, y = 5, 
             label = citation_legend,
             hjust = 0, vjust = 0,
             size = 3.25,
             color = "black")
}

# Add final labels
flux.depth.gg <- flux.depth.gg +
  labs(x = NULL, 
       y = expression("SOC sequestration (MgC ha"^-1~"year"^-1*")"),
       fill = "Sample Depth (cm)",
       subtitle = paste(data_title, normal_title, sep=""))


###### CALL THE PLOT
flux.depth.gg

##### Save the plot
ggsave(filename =  paste("SOC_flux-",normalization,"_maxdepth-",data_type,"-Depth_cols.pdf",sep=""), 
       plot = flux.depth.gg,
       width = 13, 
       height = 10.5, 
       bg = "white")
ggsave(filename =  paste("SOC_flux-",normalization,"_maxdepth-",data_type,"-Depth_cols.png",sep=""), 
       plot = flux.depth.gg,
       width = 13, 
       height = 10.5, 
       dpi = 300,
       bg = "white")


#----------------------------------------------------------------------------------------#
#--------------------- Plot colors that represent CONTINENTS ----------------------------# 

#flux.region.gg <- ggplot(data = subset(flux.df, depth_factor != "Insufficient")) +
#    #remove insufficient depth data
#    
#    # bars showing limits of flux differences by category
#    geom_col(aes(x = Cat, y = SOC_flux),
#             position = "identity", width = 0.7, fill = "gray90") +
#
#    
#    # Study-site points
#    
#    geom_point(aes(x = jittered_x, y = SOC_flux, 
#                   shape = Citation, color = Region), size = 3.5) +
#    scale_shape_manual(values = rep(15:18, 8), name = "Citation") + # cycle through shapes 15:18
#    scale_color_manual(values = region_colors) +
#    
#    # Add first three letters of citation to identify points
#    geom_text_repel(aes(x = jittered_x, y = SOC_flux,
#                        label = substr(Citation, 1, 3)), # substring of letters 1-3
#                    size = 3, segment.size = 0.3, max.overlaps = Inf) + #allow some overlap
#    
#    # Themes
#    theme_bw() +
#    theme(aspect.ratio = 0.8, text = element_text(size = 16),
#          panel.grid.major.x = element_blank(),
#          panel.grid.major.y = element_line(linewidth = .1, color = "black"),
#          legend.text =  element_text(size = 10)) +
#    geom_abline(mapping = aes(intercept = 0, slope = 0)) + 
#    coord_cartesian(xlim = c(1, 3)) +
#    
#    # control the legend order
#    guides(shape = guide_legend(order = 1), color = guide_legend(order = 2)) +
#    
#    scale_x_discrete(labels = c("No Case Control", "Cross-sectional", "Longitudinal")) +
#    
#  
#  # Add region labels
#  #     annotate("text", x = 3.9, y = lca$mean_weight_ghgs[1]+0.3, 
#  #              label = region_labels[1], color = region_colors[1], size = 3) +
#  #     annotate("text", x = 3.9, y = lca$mean_weight_ghgs[2]+0.35, 
#  #              label = region_labels[2], color = region_colors[2], size = 3) +
#  #     annotate("text", x = 3.9, y = lca$mean_weight_ghgs[3]+0.2, 
#  #              label = region_labels[3], color = region_colors[3], size = 3) +
#  #     # Add title below all rectangles
#  # annotate("text",x = 4,  y = 5, # Position below the SA rectangle
#  #          label = "Carbon Sequestration\nrequired to offset GHG\nemissions from beef",
#  #          size = 3.2,color="black",hjust = 0.5) +
#  
#    labs(x = NULL, # Categories will speak for themselves
#         y = "Difference flux of C (Mg/ha/yr)", color = "Region",
#         title = paste("Soil C sequestration - ",data_title,normal_title,sep=""))
#    
#   
#
#    
#    print(paste(data_title,";",normal_title,";","depth colors"),quote=F)
#
#ggsave(filename = paste("SOC_flux-",normalization,"_maxdepth-",data_type,"-Region_cols.pdf",sep=""), 
#       plot = flux.region.gg,
#       width = 13, 
#       height = 10.5, 
#       bg = "white")
#ggsave(filename = paste("SOC_flux-",normalization,"_maxdepth-",data_type,"-Region_cols.png",sep=""), 
#       plot = flux.region.gg,
#       width = 13, 
#       height = 10.5, 
#       dpi = 300,
#       bg = "white")

#------------------------------------------------------------------------------------------#
#####--------------------------SUMMARY TABLE-------------------------------------------#####

#Get mean, median, range, n = #, for all studies
summary <- Fluxes.study %>%
  group_by(Cat) %>% # group by category
  summarise(all.mean = round(mean(Flux.Mghayr), 2),
            all.median = round(median(Flux.Mghayr), 2),
            all.range = paste(round(min(Flux.Mghayr), 2),
                              round(max(Flux.Mghayr), 2), sep = " - "),
            all.studies = length(Flux.Mghayr))

#Get mean, med, range, n=3, for all studies excluding insufficient depth data
summary <- cbind(summary, Fluxes.study %>%
  filter(depth.tf == T) %>% 
  group_by(Cat) %>%
  summarise(exc.mean = round(mean(Flux.Mghayr), 2),
            exc.median = round(median(Flux.Mghayr), 2),
            exc.range = paste(round(min(Flux.Mghayr), 2),
                              round(max(Flux.Mghayr), 2), sep = " - "),
            exc.studies = length(Flux.Mghayr)))

summary <- summary[-6] # drop the extra cat column

# add BOOTSTRAP for med and mean CIs
bootstrap <- function(df, nboot = 1001, conf = 0.95, seed = 7012025, type = "median") {
  
  # separate out the categories
  nocc <- df %>% filter(Cat == "Time-Series Observational")
  compobs <- df %>% filter(Cat == "Comparative Observational")
  cconem <- df %>% filter(Cat == "MQE Cross-Sectional")
  ccmultm <- df %>% filter(Cat == "MQE Longitudinal")
  
  # get vectors of fluxes to sample from "nboot" times
  flux.vector.nocc <- nocc %>%
    pull(Flux.Mghayr) 
  
  flux.vector.compobs <- compobs %>%
    pull(Flux.Mghayr)
  
  flux.vector.cconem <- cconem %>%
    pull(Flux.Mghayr)
  
  flux.vector.ccmultm <- ccmultm %>%
    pull(Flux.Mghayr)
  
  # creating vectors of "nboot" NAs to be replaced in the for loops
  vector.nocc <- rep(NA, times = nboot)
  vector.compobs <- rep(NA, times = nboot)
  vector.cconem <- rep(NA, times = nboot)
  vector.ccmultm <- rep(NA, times = nboot)
  
  set.seed(seed)
  
  if (type == "median") {
    for (nn in 1:nboot) { 
      x = sample(flux.vector.nocc, replace = T, size = length(flux.vector.nocc))
      vector.nocc[nn] = median(x)
    }
    for (nn in 1:nboot) { 
      x = sample(flux.vector.compobs, replace = T, size = length(flux.vector.compobs))
      vector.compobs[nn] = median(x)
    }
    for (nn in 1:nboot) { 
      x = sample(flux.vector.cconem, replace = T, size = length(flux.vector.cconem))
      vector.cconem[nn] = median(x)
      }
    for (nn in 1:nboot) { 
      x = sample(flux.vector.ccmultm, replace = T, size = length(flux.vector.ccmultm))
      vector.ccmultm[nn] = median(x)
      }
  } else if (type == "mean") {
    for (nn in 1:nboot) { 
      x = sample(flux.vector.nocc, replace = T, size = length(flux.vector.nocc))
      vector.nocc[nn] = mean(x)
    }
    for (nn in 1:nboot) { 
      x = sample(flux.vector.compobs, replace = T, size = length(flux.vector.compobs))
      vector.compobs[nn] = mean(x)
    }
    for (nn in 1:nboot) { 
      x = sample(flux.vector.cconem, replace = T, size = length(flux.vector.cconem))
      vector.cconem[nn] = mean(x)
    }
    for (nn in 1:nboot) { 
      x = sample(flux.vector.ccmultm, replace = T, size = length(flux.vector.ccmultm))
      vector.ccmultm[nn] = mean(x)
    }
  } else {
      print("INVALID type selection. Please select type = 'median' or 'mean'.")
    }

  # get CIs
  quants <- c((1-conf)/2, 0.5, conf+(1-conf)/2) #get quants from the desired ci
  quant.nocc <- round(quantile(vector.nocc, probs = quants), 2)
  quant.compobs <- round(quantile(vector.compobs, probs = quants), 2)
  quant.cconem <- round(quantile(vector.cconem, probs = quants), 2)
  quant.ccmultm <- round(quantile(vector.ccmultm, probs = quants), 2)
  
  # categories
  category <- c("Time-Series Observational", "Comparative Observational", 
                "MQE Categorical", "MQE Longitudinal")
  
  # Create the data frame with categories as rows
  bootstrap_results <- data.frame(
    rbind(
      quant.nocc,
      quant.compobs,
      quant.cconem,
      quant.ccmultm
    )
  )

  # Rename rows
  rownames(bootstrap_results) <- category
  
  #add range 
  bootstrap_results <- cbind(bootstrap_results,
                             c(paste(bootstrap_results[1,1], bootstrap_results[1,3], sep = " - "),
                               paste(bootstrap_results[2,1], bootstrap_results[2,3], sep = " - "),
                               paste(bootstrap_results[3,1], bootstrap_results[3,3], sep = " - "),
                               paste(bootstrap_results[4,1], bootstrap_results[4,3], sep = " - ")))
  
  # Rename the columns
  colnames(bootstrap_results) <- c(paste("Q", quants[1]*100, sep = ""), 
                                   paste("Q", quants[2]*100, sep = ""),
                                   paste("Q", quants[3]*100, sep = ""),
                                   paste(conf*100, "% CI", sep = ""))
  # OUTPUT
  return(bootstrap_results)
}
# get cis
alldata.boot.med <- bootstrap(Fluxes.study, type = "median")
alldata.boot.mean <- bootstrap(Fluxes.study, type = "mean")
exc.boot.med <- bootstrap(excluded, type = "median")
exc.boot.mean <- bootstrap(excluded, type = "mean")

summary <- cbind(summary, alldata.boot.mean$`95% CI`, alldata.boot.med$`95% CI`,
                 exc.boot.mean$`95% CI`, exc.boot.med$`95% CI`)
colnames(summary) <- c("Cat", "all.mean", "all.median", "all.range", "all.studies", 
                              "exc.mean", "exc.median", "exc.range", "exc.studies",
                              "all.mean.ci", "all.med.ci", "exc.mean.ci", "exc.med.ci")

summary <- summary %>%
  select(Cat, all.studies, all.range, all.median, all.med.ci, all.mean, all.mean.ci,
         exc.studies, exc.range, exc.median, exc.med.ci, exc.mean, exc.mean.ci)


write.csv(summary, "Summary Table 1.csv", row.names = FALSE) #Table 1

#-----------------------------------------------------------------------------------------#
#------------------------------------MISC ANALYSIS-------------------------------------####
# How many of studies be animal species
Fluxes.study %>%
  group_by(Species) %>%
  summarise(Count = length(Species))

# Cross sectional studies using flux-type language
Fluxes.study %>%
  group_by(Cat) %>%
  summarise(ImpliedFlux = sum(Flux.Discussion))

# Study design over time
Fluxes.study %>%
  group_by(Cat) %>%
  summarise(mean_year = mean(as.numeric(gsub("[^0-9]", "", Year))))

# Studies by region
regional <- Fluxes.study %>%
  group_by(Region) %>%
  summarise(mean.flux = round(mean(Flux.Mghayr), 2), count = length(Flux.Mghayr),
            median.flux = round(median(Flux.Mghayr), 2), 
            range = paste(round(min(Flux.Mghayr), 2), round(max(Flux.Mghayr), 2),
                          sep = " - "))
# sites by region
regional.sites <- Fluxes %>%
  group_by(Region) %>%
  summarise(mean.flux = round(mean(Flux.Mghayr), 2), count = length(Flux.Mghayr),
            median.flux = round(median(Flux.Mghayr), 2), 
            range = paste(round(min(Flux.Mghayr), 2), round(max(Flux.Mghayr), 2),
                          sep = " - "))

# how many studies orignially reported in Mg/ha
Fluxes.study %>%
  group_by(OG.unit) %>%
  summarise(count = length(Flux.Mghayr))

# causal lang
Fluxes.study %>%
  summarise(causal = sum(causal))

#-----------------------------------------------------------------------------#
#---------------------------------Species Analysis-------------------------####
# Group by species
species.category <- rep(NA, times = length(Fluxes.study$Species))
cattle.or.mixed <- rep(NA, times = length(Fluxes.study$Species))

for (nn in 1:length(Fluxes.study$Species)) {
  

  if (is.na(Fluxes.study$Species[nn])) {
    species.category[nn] = "Not Found"

  } else if (Fluxes.study$Species[nn] == "Cattle") {
    species.category[nn] = "Cattle"
    cattle.or.mixed[nn] = "Cattle"
    
  } else if (Fluxes.study$Species[nn] == "Sheep") {
    species.category[nn] = "Small Ruminants"
    cattle.or.mixed[nn] = "Mixed"
    
  } else {
    # This covers all non-NA values that aren't "Cattle" or "Sheep"
    species.category[nn] = "Mixed Large & Small Ruminants"
    cattle.or.mixed[nn] = "Mixed"
  }
}
Fluxes.study <- cbind(Fluxes.study, species.category, cattle.or.mixed)
flux.df <- cbind(flux.df, species.category, cattle.or.mixed)
species.stats <- Fluxes.study %>%
  group_by(species.category) %>%
  summarise(mean.flux = round(mean(Flux.Mghayr), 2), count = length(Flux.Mghayr),
            median.flux = round(median(Flux.Mghayr), 2), 
            range = paste(round(min(Flux.Mghayr), 2), round(max(Flux.Mghayr), 2),
                          sep = " - "))

write.csv(species.stats, "species summary stats.csv", row.names = F)


### Mann Whitney U of cattle vs non cattle ####

#MQE Cross sectional studies only
cattle.study <- Fluxes.study %>%
  filter(cattle.or.mixed == "Cattle" & Cat == "MQE Cross-Sectional") %>%
  mutate(Flux = Flux.Mghayr) %>%
  select(Flux)

mixed.study <- Fluxes.study %>%
  filter(cattle.or.mixed == "Mixed" & Cat == "MQE Longitudinal") %>%
  mutate(Flux = Flux.Mghayr) %>%
  select(Flux)

wilcox.test(cattle.study$Flux, mixed.study$Flux)
# p = 0.08791


# Cattle, sheep, small ruminants, mixed, and none
species_colors <- c("#a6611a", "#dfc27d",  "white", "#80cdc1", "#018571")
#species_color_map <- c(
#  "Cattle" = "#a6611a", 
#  "Small Ruminants" = "#dfc27d", 
#  "Mixed Large & Small Ruminants" = "#018571", 
#  "Not Found" = "#80cdc1" )
species_shape_map <- c(
  "Cattle" = 21,
  "Small Ruminants" = 22,
  "Mixed Large & Small Ruminants" = 23,
  "Not Found" = 24)
#flux.df$species.category <- as.factor(flux.df$species.category)
#-------------------------------------------------------------------------------------#
#-------------------------------Plot with SPECIES colors---------------------------####
flux.species.gg <-
  ggplot(flux.df) +
  
 
  geom_abline(mapping = aes(intercept = 0, slope = 0)) +
  
  geom_rect(data = excl_stats,
            aes(xmin = catNum - 0.3, xmax = catNum + 0.3,
                ymin = min, ymax = max),
            fill = "gray90",
            color = NA) +  
  geom_rect(data = filter(excl_stats, catNum == 3),
            aes(xmin = catNum - 0.3, xmax = catNum + 0.3,
                ymin = q25, ymax = q75),
            fill = "gray75",
            color = NA) +
  
  geom_segment(data = excl_stats,
               aes(x = catNum - 0.3, xend = catNum + 0.3,
                   y = medn, yend = medn),
               color = "black",
               linewidth = 0.75) +
  
#  geom_point(aes(x = jittered_x, y = SOC_flux,
#                 fill = species.category), 
#             size = 5, stroke = 0.4) + 
  
  geom_point(aes(x = jittered_x, y = SOC_flux, 
                 fill = species.category, shape = species.category, color = depth.tf), 
             size = 5, stroke = 0.4) + 
  scale_fill_manual(values = species_colors, name = "Grazed Species") +
  scale_color_manual(values = c(T = "white", F = "black")) +
  scale_shape_manual(values = species_shape_map, name = "Grazed Species") +
  
  
  # Add labels
  geom_text_repel(aes(x = jittered_x, y = SOC_flux,
                      label = as.character(Label)),
                  size = 5, segment.size = 0.3,
                  max.overlaps = Inf) + 
  
  # Themes
  theme_bw() +
  theme(aspect.ratio = 0.8, text = element_text(size = 16),
        panel.grid.major.x = element_blank(),
        panel.grid.major.y = element_line(linewidth = .1, color = "black"),
        legend.text =  element_text(size = 10)) +
  
  
  
    scale_x_continuous(breaks = c(1, 2, 3, 4),
                       labels = c("Time-Series Observational", "Comparative Observational",
                                "MQE Cross-sectional", "MQE Longitudinal")) +
  
  # adding legend
  annotate("rect", 
           xmin = 3.7, ymin = 1.8, 
           xmax = 4.5, ymax = 7.5, fill = "white") +
  annotate("text", 
           x = 3.7, y = 4.67, 
           label = citation_legend,
           hjust = 0, vjust = 0.5,
           size = 3.25,
           color = "black") +
  
  
  labs(x = NULL, 
       y = expression("SOC sequestration (MgC ha"^-1~"year"^-1*")"),
       fill = "Grazed Species",
       subtitle = paste(data_title,normal_title,sep=""))
flux.species.gg
ggsave(filename =  paste("SOC_flux-",normalization,"_maxdepth-",data_type,"-Species_cols.pdf",sep=""), 
       plot = flux.species.gg,
       width = 13, 
       height = 10.5, 
       bg = "white")
ggsave(filename =  paste("SOC_flux-",normalization,"_maxdepth-",data_type,"-Species_cols.png",sep=""), 
       plot = flux.species.gg,
       width = 13, 
       height = 10.5, 
       dpi = 300,
       bg = "white")


#-------------------------------------------------------------------------------------#
#-----------------------------Non-continuous control sites-------------------------####
# Data
Sequest.nocon <- read_excel("Regenerative Grazing SOC Meta-Analysis.xlsx", 
                      sheet = "Non-continuous counterfactual")

# Rename Norm.diff.flux to Norm.diff.flux.30 to specify it's 30 cm normalization
Sequest.nocon <- Sequest.nocon %>%
    rename(Norm.diff.flux.30 = Norm.diff.flux)

#### Data Tidying & Manipulation ####
# run these all once

# Add Citation column to display author & year at once
Sequest.nocon$Citation <- paste(Sequest.nocon$Author, Sequest.nocon$Year, sep = " ")
Sequest.nocon$Citation <- factor(Sequest.nocon$Citation)
# remove the archived studies
Sequest.nocon <- Sequest.nocon %>%
    filter(Archive == FALSE)

# rename all continents besides NAm, SAm, and Aus to "Other" and relevel
Sequest.nocon <- Sequest.nocon %>%
    mutate(
        Region = Region %>%
            #  collapse all other continents into "Other"
            forcats::fct_other( keep = c("North America", "South America", "Australia"),
                                other_level = "Other") %>%
            forcats::fct_relevel("North America", "South America", "Australia", "Other")
    )

# Pull out ONLY difference fluxes
nocon <- Sequest.nocon %>%
    filter(Difference.flux.tf == TRUE)


# Reorder levels of Cat to change column order
nocon$Cat <- factor(nocon$Cat, 
                    levels = c("Time-Series Observational", "Comparative Observational", 
                               "MQE Cross-Sectional", "MQE Longitudinal"))

# sum all depth slices and round to nearest depth level specified above
nocon <- nocon %>%
    mutate(Sample.Depth.cm = select(., starts_with("Slice")) %>%
               rowSums(na.rm = TRUE)) %>%
    mutate(Sample.Depth.cm = map_dbl(Sample.Depth.cm,
                                     ~depth_levels[which.min(abs(depth_levels - .x))]))

# create a column that counts depth slices
nocon <- nocon %>%
    mutate(Slices = str_count(All_depths.cm, ",") + 1)

# Create a jittered x-value column to have predictable jitter
set.seed(07072025) 
nocon$jittered_x <- jitter(as.numeric(nocon$Cat), amount = 0.3)

# Create a binary variable for robust depth info
nocon$depth.tf <- ifelse(nocon$Sample.Depth.cm > 10 & 
                             nocon$Slices > 1, TRUE, FALSE)

# Allow for custom coloring later
nocon$depth_factor <- factor(
    ifelse(nocon$depth.tf == FALSE, "Insufficient", as.character(nocon$Sample.Depth.cm)),
    levels = c("20", "30", "50", "100", "Insufficient"))


unique_citations <- nocon %>%
  group_by(Citation) %>%
  summarise(Citation = first(Citation)) # get all unique citations

# giving each unique citation a number 
unique_citations$Label <- c(as.character(1:(length(unique_citations$Citation))))


# left-join by shared "Citation". result should work for site and study wise plots
nocon <- nocon %>% 
  left_join(unique_citations, by = "Citation")

nocon <- nocon %>% # adding a column for a legend
  mutate(citation_display = paste0(Citation, " - ", Label)) %>%
  # Create a sorting key based on Label 
  mutate(label_numeric = as.numeric(gsub("[a-z]", "", Label)),
         
         label_letter = gsub("[0-9]", "", Label)) %>%
  
  arrange(label_numeric, label_letter) %>%
  
  mutate(citation_display = factor(citation_display, levels = unique(citation_display))) %>%

  select(-label_numeric, -label_letter)

nocon$noncont_cat <- nocon$nc_control.cat

nocon.study <- nocon %>%
  # Create a df of study-wise difference fluxes
  group_by(Citation, noncont_cat) %>%
  # using first() to retain necessary columns from original df
  summarize(Author = first(Author), Year = first(Year), Cat = first(Cat), 
            All_depths.cm = first(All_depths.cm), Sample.Depth.cm = first(Sample.Depth.cm),
            Slices = first(Slices), jittered_x = first(jittered_x), depth.tf = first(depth.tf),
            depth_factor = first(depth_factor), Significant.tf = first(Significant.tf),
            Region = first(Region), Significance_tested.tf = first(Significance_tested.tf),
            Species = first(Species), Flux.Discussion = first(Flux.Discussion),
            Label = first(Label), citation_display = first(citation_display), 
            OG.unit = first(OG.unit), causal = first(Causal.Language),
            # mean flux for each study
            Flux.Mghayr = mean(Flux.Mghayr),
            Norm.diff.flux.30 = mean(Norm.diff.flux.30),
            Norm.diff.flux.100 = mean(Norm.diff.flux.100))

citation_legend <- nocon %>%
  select(Label, Citation) %>%
  distinct() %>%
  arrange(as.numeric(gsub("[a-z]", "", Label))) %>%  # Sort numerically
  mutate(legend_text = paste0(Label, ": ", Citation)) %>%
  pull(legend_text) %>%
  paste(collapse = "\n")

set.seed(555)
nocon$jittered_x <- jitter(as.numeric(nocon$Cat), amount = 0.3)
#----------------------------------------------------------------------------------#
####----------------------Plots with NON CONT control---------------------------####
nocon.sites.gg <- 
ggplot(nocon) +
  #reinforce zero line
  geom_abline(aes(slope = 0, intercept = 0), color = "gray5") +
  
  #points
  geom_point(aes(x = jittered_x, y = Flux.Mghayr, fill = depth.tf, shape = noncont_cat),
             size = 5, color = "black", stroke = 0.3) +
  #point labels
  geom_text_repel(aes(x = jittered_x, y = Flux.Mghayr, label = Label),
                  max.overlaps = Inf) +
  
  #Points with insuff depth are whited out
  scale_fill_manual(values = c("FALSE" = "white", "TRUE" = "gray15"),
                    name = "Non-Continuous Control") +
  
  #Category of non-cont control group
  scale_shape_manual(values = c("Exclusion" = 22, "Fire" = 23, "Hay Harvest" = 24,
                                "Wild Grazing" = 25, "Relaxed Rotation" = 21), 
                     name = "Non-Continuous Control") +
  
  #Crop out the time-series because n=0
  scale_x_continuous(breaks = c(2, 3, 4),
                     labels = c("Comparative Observational",
                                "MQE Cross-sectional", "MQE Longitudinal")) +
  
  #Adding y-axis ticks
  scale_y_continuous(breaks = c(-5, -2.5, 0, 2.5, 5, 7.5)) +
  
  guides(fill = "none") + # remove the fill legend
  
  # Labels
  annotate("rect", 
           xmin = 3.9, ymin = 3.8, 
           xmax = 4.5, ymax = 7.49, 
           fill = "white") +
  #text
  annotate("text", 
           x = 4, y = 4.6, 
           label = citation_legend,
           hjust = 0, vjust = 0,
           size = 3.25,
           color = "black") +
  
  #Themes
  theme_bw() +
  theme(text = element_text(size = 16),
        panel.grid.major.x = element_blank(),
        panel.grid.major.y = element_line(linewidth = .1, color = "black"),
        legend.text =  element_text(size = 10)) +
  
  #Labels and titles
  labs(x = NULL, 
       y = expression("SOC sequestration (MgC ha"^-1~"year"^-1*")"),
       title = "One point per pasture site")


nocon.studies.gg <- 
  ggplot(nocon.study) +
  #reinforce zero line
  geom_abline(aes(slope = 0, intercept = 0), color = "gray5") +
  
  #points
  geom_point(aes(x = jittered_x, y = Flux.Mghayr, fill = depth.tf, shape = noncont_cat),
             size = 5, color = "black", stroke = 0.3) +
  #point labels
  geom_text_repel(aes(x = jittered_x, y = Flux.Mghayr, label = Label),
                  max.overlaps = Inf) +
  
  #Points with insuff depth are whited out
  scale_fill_manual(values = c("FALSE" = "white", "TRUE" = "gray15"),
                    name = "Non-Continuous Control") +
  
  #Category of non-cont control group
  scale_shape_manual(values = c("Exclusion" = 22, "Fire" = 23, "Hay Harvest" = 24,
                                "Wild Grazing" = 25, "Relaxed Rotation" = 21), 
                     name = "Non-Continuous Control") +
  
  #Crop out the time-series because there are none
  scale_x_continuous(breaks = c(2, 3, 4),
                     labels = c("Comparative Observational",
                                "MQE Cross-sectional", "MQE Longitudinal")) +
  
  #Adding y-axis ticks because default is + or - 5
  scale_y_continuous(breaks = c(-5, -2.5, 0, 2.5, 5, 7.5)) +
  
  guides(fill = "none") + # remove the fill legend
  
  # Labels
  annotate("rect", 
           xmin = 3.9, ymin = 0.1, 
           xmax = 4.5, ymax = 2.0, 
           fill = "white") +
  #text
  annotate("text", 
           x = 4, y = 0.5, 
           label = citation_legend,
           hjust = 0, vjust = 0,
           size = 3.25,
           color = "black") +
  
  #Themes
  theme_bw() +
  theme(text = element_text(size = 16),
        panel.grid.major.x = element_blank(),
        panel.grid.major.y = element_line(linewidth = .1, color = "black"),
        legend.text =  element_text(size = 10)) +
  
  #Labels and titles
  labs(x = NULL, 
       y = expression("SOC sequestration (MgC ha"^-1~"year"^-1*")"),
       title = "One point per study")


#Sites nocon
ggsave(filename =  paste("nocon sites.png",sep=""), 
       plot = nocon.sites.gg,
       width = 13, 
       height = 10.5, 
       bg = "white")

#studies nocon
ggsave(filename =  paste("nocon studies.png",sep=""), 
       plot = nocon.studies.gg,
       width = 13, 
       height = 10.5, 
       bg = "white")


nocon.summary <- nocon.study %>%
  group_by(Cat) %>%
  summarise(studies = length(Flux.Mghayr),
            median = round(median(Flux.Mghayr), 2),
            mean = round(mean(Flux.Mghayr), 2),
            range = paste(round(min(Flux.Mghayr), 2), 
                          round(max(Flux.Mghayr), 2), sep = " - "))
```
